# A functional Bayesian model for hydrogen-deuterium exchange mass-spectrometry

**DOI:** 10.1101/2022.07.18.500413

**Authors:** Oliver M. Crook, Chun-wa Chung, Charlotte M. Deane

## Abstract

Proteins often undergo structural perturbations upon binding to other proteins or ligands or when they are subject to environmental changes. Hydrogen deuterium exchange mass-spectrometry (HDX-MS) can be used to explore conformational changes in proteins by examining differences in the rate of deuterium incorporation in different contexts. To determine deuterium incorporation rates, HDX-MS measurements are typically made over a time course. Recently introduced methods show that incorporating the temporal dimension into the statistical analysis improves power and interpretation. However, these approaches have technical assumptions which hinder their flexibility. Here, we propose a more flexible methodology by reframing these methods in a Bayesian framework. Our proposed framework has improved algorithmic stability, allows us to perform uncertainty quantification, and can calculate statistical quantities that are inaccessible to other approaches. We demonstrate the general applicability of the method by showing it can perform rigorous model selection on a spike-in HDX-MS experiment and improved interpretation in an epitope mapping experiment. Bayesian analysis of an HDX experiment with an antibody dimer bound to an E3 ubiquitin ligase identifies at least two interaction interfaces where previous methods obtained confounding results due to the complexities of conformation change on binding. Our findings are consistent with the co-crystal structure of these proteins, demonstrating a bayesian approach can identify important binding epitopes from HDX data.

## 1 Introduction

Protein structures can be perturbed due to alterations in their context and HDX-MS is a powerful technique to examine these perturbations (Orengo *et al*., 1999; Engen, 2009; Chalmers *et al*., 2011; Houde *et al*., 2011; Masson *et al*., 2017; Sauve *et al*., 2018). When a protein is incubated in heavy water its amide hydrogens exchange with deuterium at a context-specific rate (Katta and Chait, 1993). This rate of exchange will be in accordance with Linderstrom-Lang theory (James *et al*., 2021) and is also affected by solvent accessibility, topological flexibility, amino-acid content, secondary structure and conformal heterogeneity (Konermann *et al*., 2011; Kammari and Topp, 2020; Jia *et al*., 2020). Since deuterium is heavier than hydrogen, we can measure rates of incorporation by examining mass-shifts and isotopic expansion using bottom-up mass-spectrometry (Masson *et al*., 2017). To mediate complex protein kinetics HDX-MS is performed over a time-course, examining the deuterium incorporated at different exposure times of the sample to heavy water (Crook *et al*., 2022b). We recently showed that using tools from functional data analysis and empirical Bayes analysis could increase power, reduce false positives and improve interpretation of HDX-MS experiments when compared to t-test and linear mixed models (Crook *et al*., 2022b). Furthermore, that functional data analysis approach could be applied to large epitope mapping experiments where there were no previously rigorous statistical tools. This approach fits either logistic or Weibull kinetics to HDX-MS data, then, to assess the quality of fits, an F-statistic is computed. This statistic is then *moderated* using an empirical Bayes approach (Crook *et al*., 2022b).

However, to our earlier approach (Crook *et al*., 2022b) required restrictive assumptions on the kinetics which reduces power. Here, we recast our analysis in the Bayesian framework which allows us to further increase the flexibility of the model (Gelman *et al*., 2013). Bayesian analysis has a number of further advantages (Gelman *et al*., 2013). First, it allows us to shrink residuals towards zero, meaning that differences between HDX kinetics are easier to identify. Secondly, it allows us to regularise the inferred parameters allowing us to fit functional models with more parameters than observations. Thirdly, model fitting is less sensitive to initialisation and so produces more reliable results (Gelman *et al*., 2013). In addition, Bayesian analysis allows us to quantify uncertainty using probability distributions, allowing us to report the confidence in our results (Betancourt, 2021). This also allows us to compute quantities that are impossible to obtain without using a Bayesian approach, for example the probability that a deuterium difference exceed a particular value. A Bayesian method for HDX-MS data has previously been developed Saltzberg *et al*. (2017), but the focus of that work is on determining residue-level information rather than statistical differences at the peptide level. Furthermore, that approach employs clustering before significance testing, which means the hypothesis tested is conditioned on the data, which violates selective inference protocols and hence inflates false positives (Taylor and Tibshirani, 2015).

First, we show that our approach is consistent: controlling false positives and obtaining true positives in simulations. We then examine competing models for HDX and show that when there are EX1 dynamics, a Weibull model is preferred over a logistic model for HDX data. We characterise this in a spike-in HDX experiment showing a strong preference for almost all peptides to follow Weibull-type kinetics (Hageman and Weis, 2019). However, we find no evidence to support a functional mixed-model for HDX. Finally, we perform epitope mapping in HOIP-RBR (Tsai *et al*., 2020) showing that we can increase the number of findings in these experiments when compared to an empirical Bayes approach (Crook *et al*., 2022b). In particular, we were previously unable to identify the binding epitope of dAb3 from the HDX data using any previous statistical method; however, our Bayesian analysis of this HDX experiment suggests three possible binding interfaces which are supported by the co-crystal structure of the dAb3 dimer and HOIP-RBR. In this case, we present an array of new visualisations and computational techniques afforded by taking a probabilistic approach.

## 2 Main

### 2.1 Model Summary

We begin with a narrative description of our model, with technical details given in the methods. We propose to model HDX-MS data using functional models (time-dependent curves). We choose to use smooth parametric models for the modelling, either using a logistic or Weibull-type model, depending on the expected kinetics. We then place prior distributions on the parameters of these models and Bayes’ theorem tells us that after observing data we can update these prior distributions into posterior distributions (Gelman *et al*., 2013). These posterior distributions quantify the uncertainty in the model parameters. The Bayesian framework allows us to compute probability distributions as a function of time, as well as the probability of a model given the data. For example, we can compute whether covariate dependent models are preferred over models that are blinded to the covariates. Covariates in this scenario would be any conditions, states or general quantities that could potentially perturb HDX kinetics. This preference is quantified through the posterior probability of that particular model. In this manuscript, we introduce several new quantities, such as the probability of deuterium incorporation being above a certain value. These quantities are difficult to compute in other statistical frameworks. In addition, the prior distribution acts as regularisation for our model parameters and hence rules-out improbable parameters - this stabilises the inference in our model. For a review of Bayesian methods applied to proteomics data, we refer to Crook *et al*. (2022a).

### 2.2 Simulations show that the Bayesian approach is powerful and calibrated

First, we characterise the performance of our approach in simulation scenarios (see methods), which cover a range of expected situations in HDX studies. We perform simulations with four or five time points and two or three replicates.

We examine two quantities whilst performing these simulations. The first is to compare performance to our previously proposed empirical Bayes method for HDX-MS data (Crook *et al*., 2022b) (which was shown to be the most powerful method for determining deuterium differences in HDX-MS data). Here, we use the area-under-the-curve (AUC) of the TPR (True Positive Rate) - FPR (False Positive Rate) curve. Values close to 1 indicate perfect performance, whilst values of 0.5 suggest random guessing. We run five simulations in each setting and report the distribution. We find that both methods give AUC values close to 1 (see Figure 1). The minor differences between the two methods are too small to be relevant in practice.

**Figure 1:**
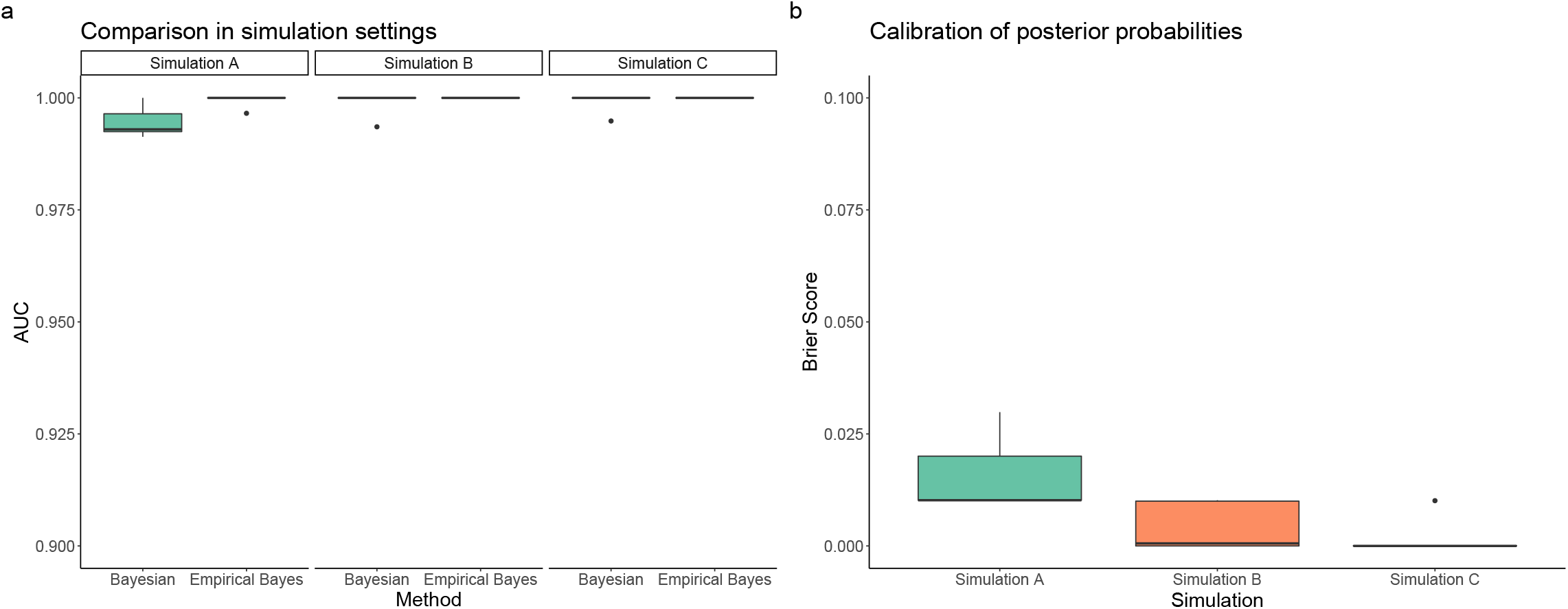
Simulation study to compare statistical methods on HDX-MS data. (a) Area-under-the-curve (AUC) for both the proposed Bayesian approach and the empirical Bayesian approach. The effective difference are too small to suggest any practical difference between the approaches (b) Brier scores for our Bayesian approach. Values are close to 0 suggesting high quality calibration. See methods for details on simulations; briefly, simulation A has four time points and three replicates. Simulation B has four time points and two replicates. Simulation C has five time points and two replicates.

In addition, we compute the Brier score (see methods) for our Bayesian approach. The Brier score assesses whether the posterior probabilities are correctly calibrated. This means that if there is a perturbation to the HDX kinetics, we expect probabilities close to 1 and if there is no perturbation, we expect probabilities close to 0. The Brier score quantifies the average calibration-error. As can be seen from Figure 1 the errors are, on average, less than 1 percentage point. This means that if there are no perturbations to the HDX-MS kinetics our approach could report a probability of 0.01 of a perturbation and if there is a perturbation our approach could report a probability of 0.99 of a perturbation. This represents almost perfect calibration and our probabilities can be interpreted as forecasts. Given that our Bayesian approach has similar statistical performance to the empirical Bayes approach, the remainder of the manuscript focuses on demonstrating the interpretation advantages of our Bayesian method.

### 2.3 Bayesian model selection for HDX

#### 2.3.1 A Weibull model for EX1 kinetics

When a portion of amides undergo correlated deuterium exchange due to simultaneous unfolding and refolding, we observe EX1 kinetics. EX1 kinetics describe different populations of proteins undergoing different hydrogen-deuterium exchange mechanisms. This results in multi-modal deuterated spectra, in which each mode corresponds to a different protein subpopulation. This behaviour can be challenging to model. Briefly, at the level of each residue a logistic model is an appropriate model of exchange:

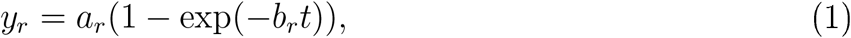

where *r* indexes residues, *b*_*r*_ denotes the rate constant, *a*_*r*_ the deuterium recovery and *t* is the time of exposure to heavy water. However bottom-up mass-spectrometry measures peptides and so the observation process is:

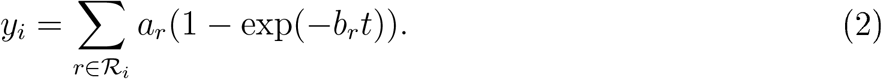

Here *i* index peptides and ℛ_*i*_ the set of exchangeable residues for peptide *i*. However, in practice, it is challenging to fit this model because of the number of parameters typically exceeds the number of observations. Therefore, it is required to approximate the kinetics for each peptide. Some suggested models include the following (Chetty *et al*., 2009; Crook *et al*., 2022b):

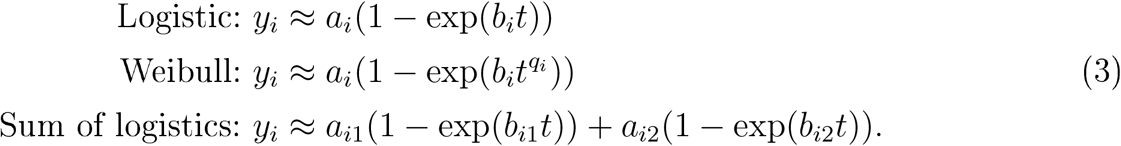

A more complex possibility is a non-parametric model *y*_*i*_ ≈ *f*(*t*), but we have chosen to explore if the simpler models are sufficient to explain the majority of HDX-MS kinetics. Currently, there is no statistical methodology or guidance on determining which approximation to use for statistical testing for peptide-centric HDX-MS data. We are particularly interested in determining the preferred model for data with EX1 kinetics. Here, using Bayesian statistics, we test whether a logistic or Weibull model is a better model of centroided EX1 kinetics (see methods). To perform this analysis, we simulate EX1 kinetics for 100 peptides. The length of each peptide was sampled uniformly between 5 and 15, with the amino acids chosen uniformly at random. The first mode was assumed to have 0 deuterium incorporation, whilst the incorporation for the second mode was sampled uniformly from [0, 1] and the charge state was sampled uniformly between one and eight. Deuterium incorporations measurements were made at 0, 300, 500, 700 and 1000 seconds post exposure to heavy water. The relative proportion of the second mode was assumed to rise exponentially. For each peptide two replicates were simulated to allow for natural variations. Bimodal spectra were simulated using the natural isotope distributions for that peptide and the centroid computed. To examine whether a logistic or Weibull model was preferred, we used the posterior probability of each model, as well as the leave-one-out expected log predictive density - a measure of out of sample predictive performance of our models (see methods for more details). Figure 2 plots paired boxplots which show that the Weibull model was preferred according to both measures. This is despite the additional penalisation of the Weibull model because of a more diffuse prior.

**Figure 2:**
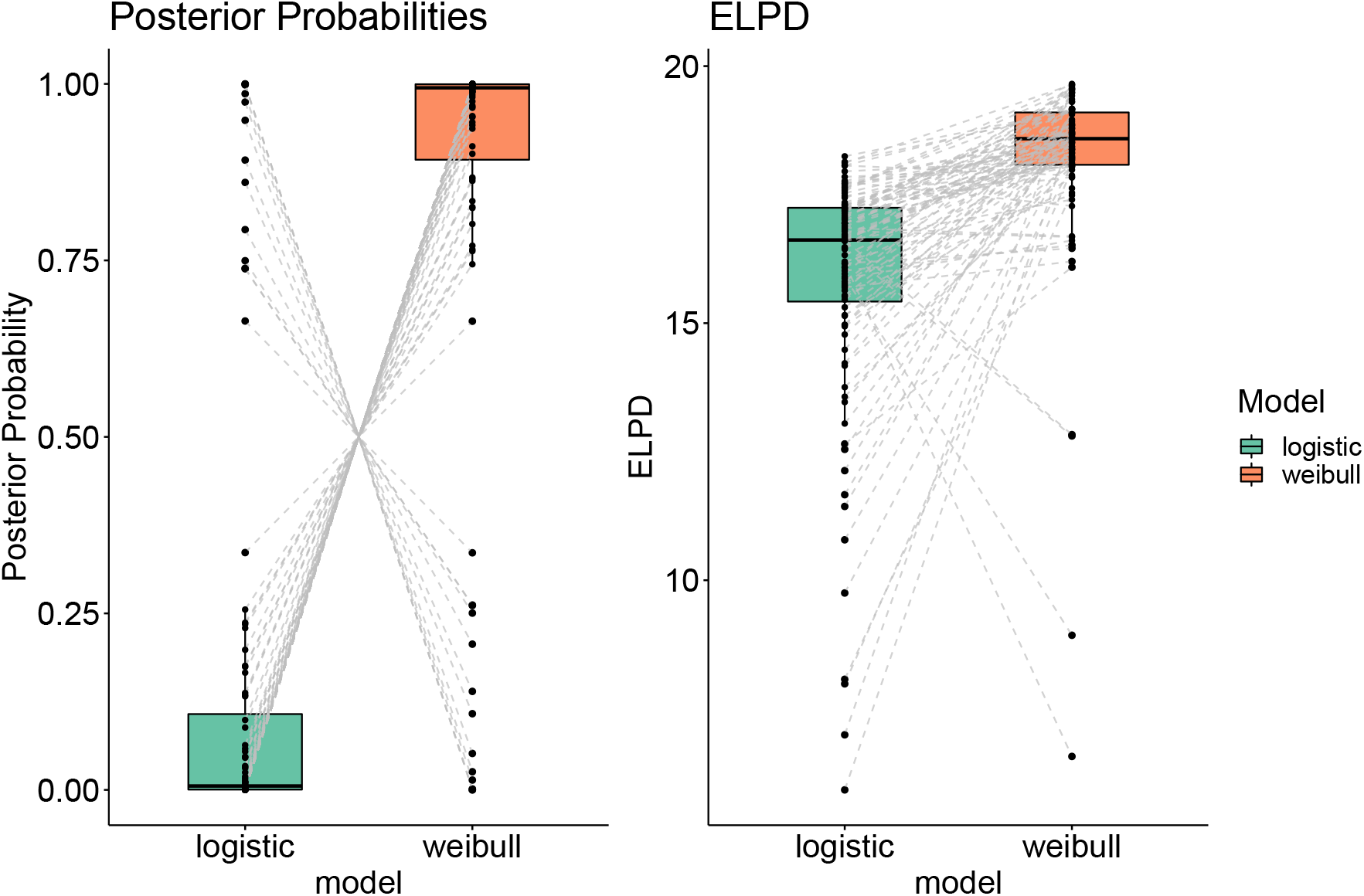
Examining model selection for EX1 kinetics. Paired boxplots for the two metrics of interest: posterior model probabilities (left) and leave-one-out expected log predictive density (ELPD) (right). Grey lines indicating the simulation pairings, indicating not only preference on average for the Weibull model but the vast majority of peptides.

#### 2.3.2 Model selection for structural spike-in experiment

Having analysed simulated data, we next analyse a structural spike-in experiment, where HDX data on maltose-binding protein (MBP) was generated in seven replicates across four HDX labelling times (Hageman and Weis, 2019). Additional experiments were carried out in triplicate for the W169G (tryptophan residue 169 to glycine) structural variant. Here MBP-W169G was spiked into the wild-type MBP sample in 5, 10, 15, 20, 25% proportions, and a further experiment included a 100% mutant sample. All data were analysed on a Agilent 6530 Q-TOF mass spectrometer and raw spectra processed in HDExaminer. Since there are two populations of proteins (either the WT or W169G), these HDX data undergo multi-modal exchange dynamics. This allows us to further test our previous model selection approach in practice. Furthermore, as these experiments are replicated we can also examine a random-effects model which could account for random variations in the plateauing of the HDX kinetics (see methods) across replicates. Thus, there are now three possible models to consider: a logistic model, a Weibull model, a Weibull model with random plateaus. To compare these models formally, we again use posterior model probabilities and the leave-one-out expected log predictive density (ELPD). For brevity, we consider the 10% and 15% spike-in experiments.

Figure 3 shows ternary plots for these three models, where the metrics have been rescaled to indicate relative model preference. Each pointer represents a modelled peptide. In general, the posterior probability suggests a preference for the Weibull model. Occasionally, there is a preference for logistic model with little support for the more complex random plateaus model. The conclusions are somewhat similar when using ELPD, but the random plateau model has a little more support. This suggests that the preferred model is a Weibull model and the more complex random plateau model is unnecessarily complex but might be useful in predicting new data. Hence, for under-determined HDX-MS data, where there are fewer measured data points than the full kinetic model, Bayesian modelling offers a principled approach to determine an approximate model. In the supplementary material, we also show that our Bayesian approach does not generate false positives in a null permutation experiment.

**Figure 3:**
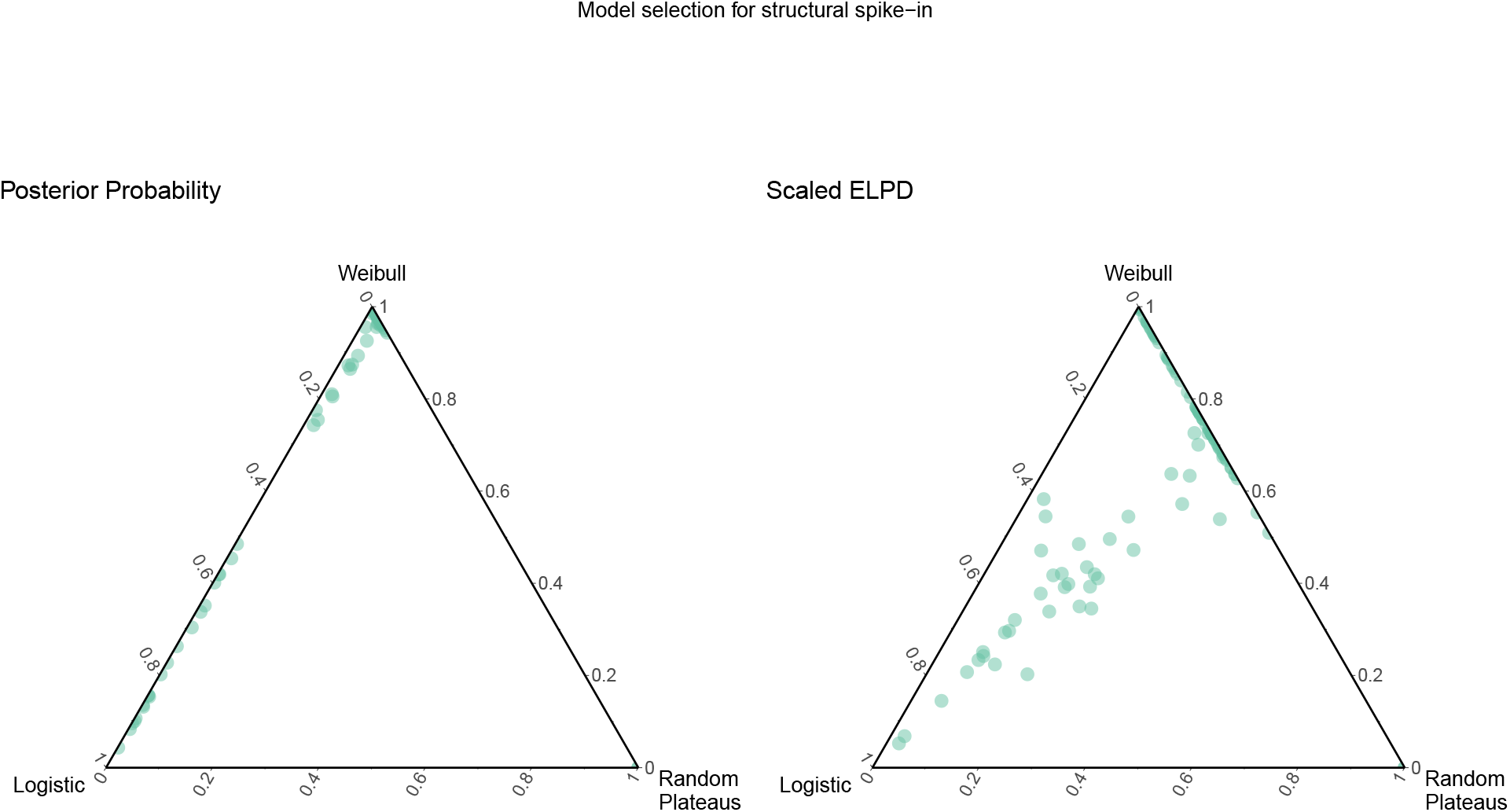
Examining model selection for structural spike-in kinetics. Ternary plots for the two metrics of interest: posterior model probability (left) and ELPD_LOO_ (right). We consider 3 models: a Logistic model (left), a Weibull model (top) and Random Pleateus (right). The metrics have been rescaled to indicate relative model preference. Each pointer represent the metrics for a modelled peptide. The closer the pointer is to the apex indicating the model, the greater the preference for that model.

### 2.4 Case study: Epitope mapping of HOIP-RBR

Ubiquitination is a key post-translational modification and acts as a molecular toggle in the regulation of cellular processes. HOIP is a member of the E3 complex LUBAC and plays a significant role in immune signalling, by conjugating linear polyubiquitin chains (Kirisako *et al*., 2006; Gerlach *et al*., 2011; Spratt *et al*., 2014). The active domain of the LUBAC complex is located in the HOIP-RBR domain and inhibition of LUBAC activity has been shown to be of potential therapeutic benefit. To support structure-based inhibitor design, Tsai *et al*. (2020) sought to develop single chain antibody-based crystallization chaperones. As part of their study, they performed epitope mapping of a library of synthetic domain antibodies (dAbs) using HDX-MS. To identify binding epitopes, they looked for “protection” signatures, that is surface amides that incorporate deuterium more slowly in the bound state as they are shielded from the solvent. Tsai *et al*. (2020) performed HDX-MS experiments for HOIP-RBR upon single domain antibody (dAb) complexation and in the APO state. Massspectrometry was performed using a Waters Synapt G2-Si instrument and raw data was processed using DynamX. HDX-MS measurements were taken at 0, 30 and 300 seconds post exposure to heavy water, for thirteen dAbs at different molar concentrations. Only a single replicate measurement was taken in each state so that measurements of many different dAbs could be made. This means that statistically rigorous analysis is challenging and only obvious binding epitopes could be identified (Tsai *et al*., 2020; Crook *et al*., 2022b). Here, we apply our Bayesian model to two of the dAbs with the hope of quantifying our confidences in the binding epitope and the possible conformational changes of HOIP upon dAb-complexation.

#### 2.4.1 HDX-MS analysis of HOIP-RBR-dAb25

First, we focus on dAb25, one of 13 dAbs, that demonstrated non-standard HDX behaviour for this complex (Tsai *et al*., 2020; Crook *et al*., 2022b). By visual examination, Tsai *et al*. (2020) noted deprotection signatures and, thus, hypothesised that dAb25 locks HOIP-RBR in a more open conformation. However, this qualitative approach is prone to bias and errors. They could not apply a quantitative approach such as the t-test or a linear mixed model because neither of these were applicable to the data due to lack of replication. Recently, an empirical Bayes function model was able to identify statistical signficant differences (Crook *et al*., 2022b). That analysis identified eight peptides for which the deuterium kinetic were altered (adjusted *p*-value < 0.05), suggesting that the binding epitope was located in the inbetween ring (IBR) of HOIP (see supplementary figure 14). However, that approach required the fixing of the parameters in a Weibull model to *b* = 0.5, *p =* 1, *d =* 0. This reduces the flexibility of the model and hence kinetics that deviate from these assumptions will be poorly modelled. Our Bayesian approach allows us to regularise rather than fix these parameters facilitating modelling of a larger range of kinetics. In particular, the rate constant *b* can be inferred rather than fixed. Our proposed Bayesian logistic model is fitted using Markov-chain Monte-Carlo (MCMC) and so uncertainty in these parameters is propagated to the uncertainty in the underlying time-dependent function.

Our Bayesian approach facilitates a number of probabilistic computations. First, we examine cases where there is increased deuterium exchange as a result of antibody binding. That is regions of HOIP-RBR that are more solvent accessible. For this, we compute the 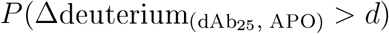 for fixed values of *d*, where we use Δdeuterium_(*A,B*)_ to denote the deuterium difference between states *A* and *B*. This computation was made at 300 seconds post exposure to heavy water with time dependent computations made later. Figure 4a shows a plot of this probability as a function of peptide, arranged in protein order. First, we observe that peptides generally appear to increase their deuterium incorporation when the antibody is bound, suggesting that this antibody holds HOIP-RBR in a more open conformation. Secondly, we note that these effect sizes are generally small, suggesting that these changes are likely subtle. Finally, we observe a clear region in the centre of the plot where deuterium incorporation appears to be reduced, suggesting a likely binding epitope. To examine whether these kinetics are consistent with respect to the time dimension of the HDX measurements, we can compute 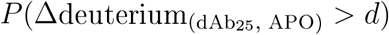 for a fixed *d* = 0.5 for each time point. Figure 4b plots these quantities for each peptide. This plot supports the same conclusions as Figure 4a.

**Figure 4:**
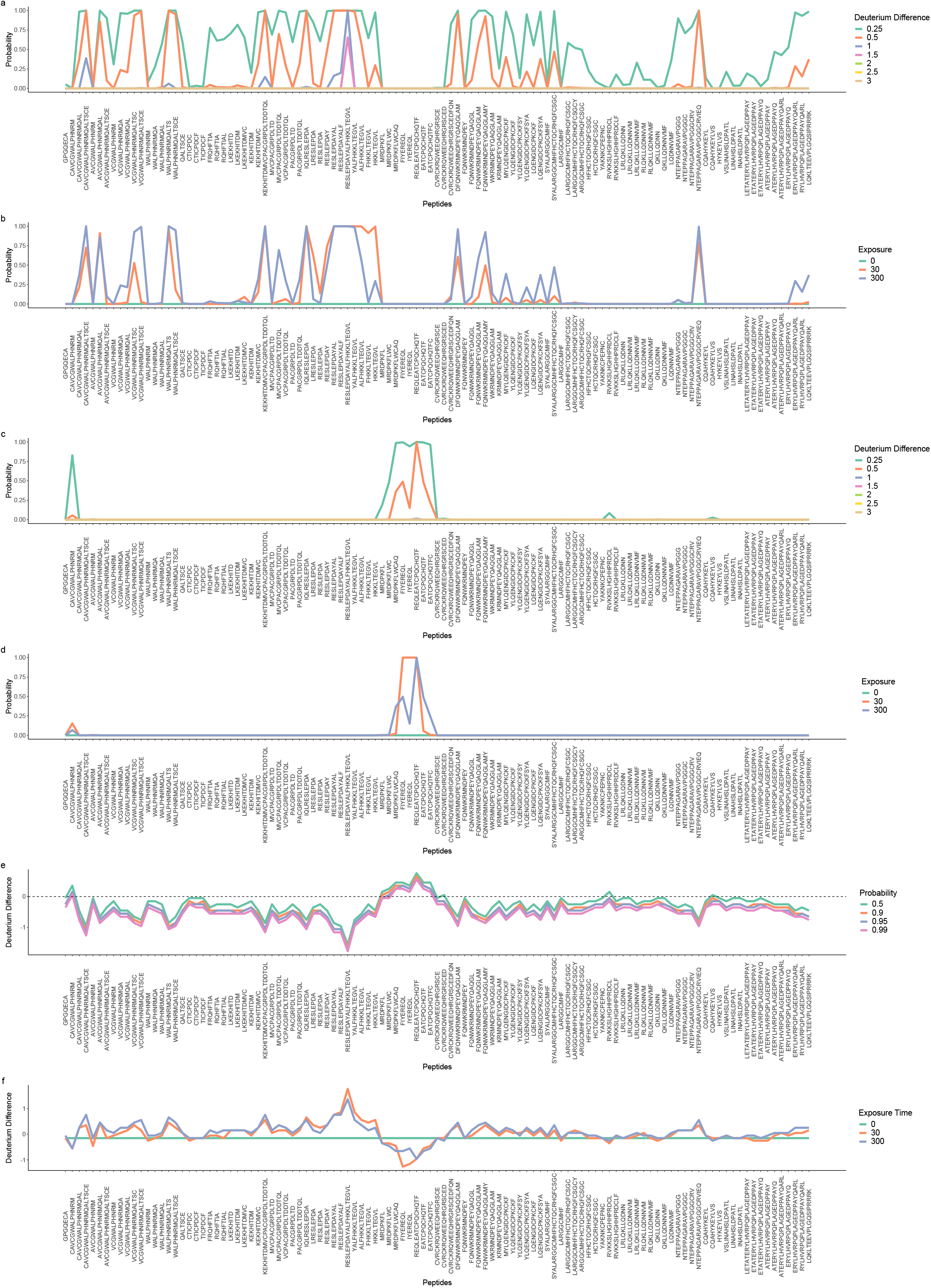
Posterior quantities of interest from Bayesian analysis. (a) The probability that the deuterium difference is higher than a particular effect size in the dAb-bound state, in general we see a tendency for deuterium to be incorporated in the bound state. Though a clear region of no deuterium incorporation is visible (b) Consistency of probabilities across the temporal dimension of the data. Plotted is the probability that the deuterium difference is higher than 0.5 in the dAb-bound state. (c) The probability that the deuterium difference is greater in the apo state (epitope mapping) for a number of effect sizes. An epitope is clearly visible. (d) Consistency of probabilities across the temporal dimension of the data. Plotted is the probability that the deuterium difference is greater in the apo state. An identified epitope is consistent across the temporal dimension. (e) We plot the largest value *d* such that *P*(Δ*D* > *d*) = *p* for different values of *p*, the dAb-bound state is used as the reference. We see that in general a tendency of increase incorporation in the dAb-bound state and the epitope is clearly identifiable (cluster of values above 0). (f) Consistency across the temporal dimension of the data. We plot the largest value *d* such that *P*(Δ*D* > *d*) = p for different values of *p*, the apo state is used as the reference. We see a general tendency for increased incorporation of deuterium in the dAb-bound state. The epitope is identifiable from the cluster of values below 0.

To identify the epitope, we switch the ordering in our computation 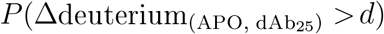. We compute the relevant quantities again and plot in Figures 4c and d. Here, the probable epitope is evident from a cluster of several peptides in the centre of the figures. This includes a cluster of seven peptides with some probability of reduced deuterium exchange. This more than triples the previous evidence for an epitope in this location given by the empirical Bayes approach. This suggests that we have confidently identified the epitope rather than spurious measurement fluctuations. Since our approach is probabilistic, these probabilities could be used alongside crystal structure data or machine learning based approaches as complementary evidence.

More elaborate probabilistic computations are possible. For example, we can consider the equation 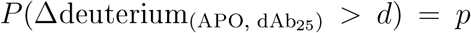 for some *p*, say 0.99. We note that if *d*_1_ > *d*_2_ then 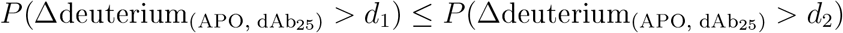 because the integration is over a larger set. Thus decreasing the value of *d*, cannot decrease *p*. From here, we can find the largest value *d* such that 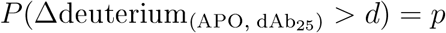. We plot this quantity for different values of *p* in Figure 4e. Of course, if the largest such value of *d* is less than zero, this suggests that deuterium is being incorporated in those peptides in the antibody bound state. This procedure allows us to clearly identify the threshold at which peptides become interesting and facilitates identification of the peptide with the strongest evidence for differences. This plot further highlights the probable binding epitope. For illustration, we include the temporal version of this plot in Figure 4f, now taking the apo state as the reference and letting *p* = 0.95.

Finally, we examine a peptide in the RING1 region of HOIP-RBR (MVCPACGRPDLT-DDTQL [742-758], see supplementary figure 14 b), which has an increase in deuterium incorporation upon dAb25 complexation. We can visualise a number of posterior quantities from this analysis, including the posterior predictive distribution of the entire time resolved kinetics (Figure 5a); the posterior predictive distribution, as a violin plot, of deuterium incorporation at specific times (Figure 5b) and the posterior predictive distribution, as a violin plot, of the deuterium difference at specific times (Figure 5c). These plots allow us to see that a deuterium difference is already evident by 30 seconds but is much more pronounced at 300 seconds. We are able to report probabilities and hence confidence in our conclusions, allowing us to weigh up HDX-MS data in the context of other experiments. In summary, our Bayesian approach allows powerful visualisation and computation for HDX data and increases our ability to carefully draw quantitative insights from our data.

**Figure 5:**
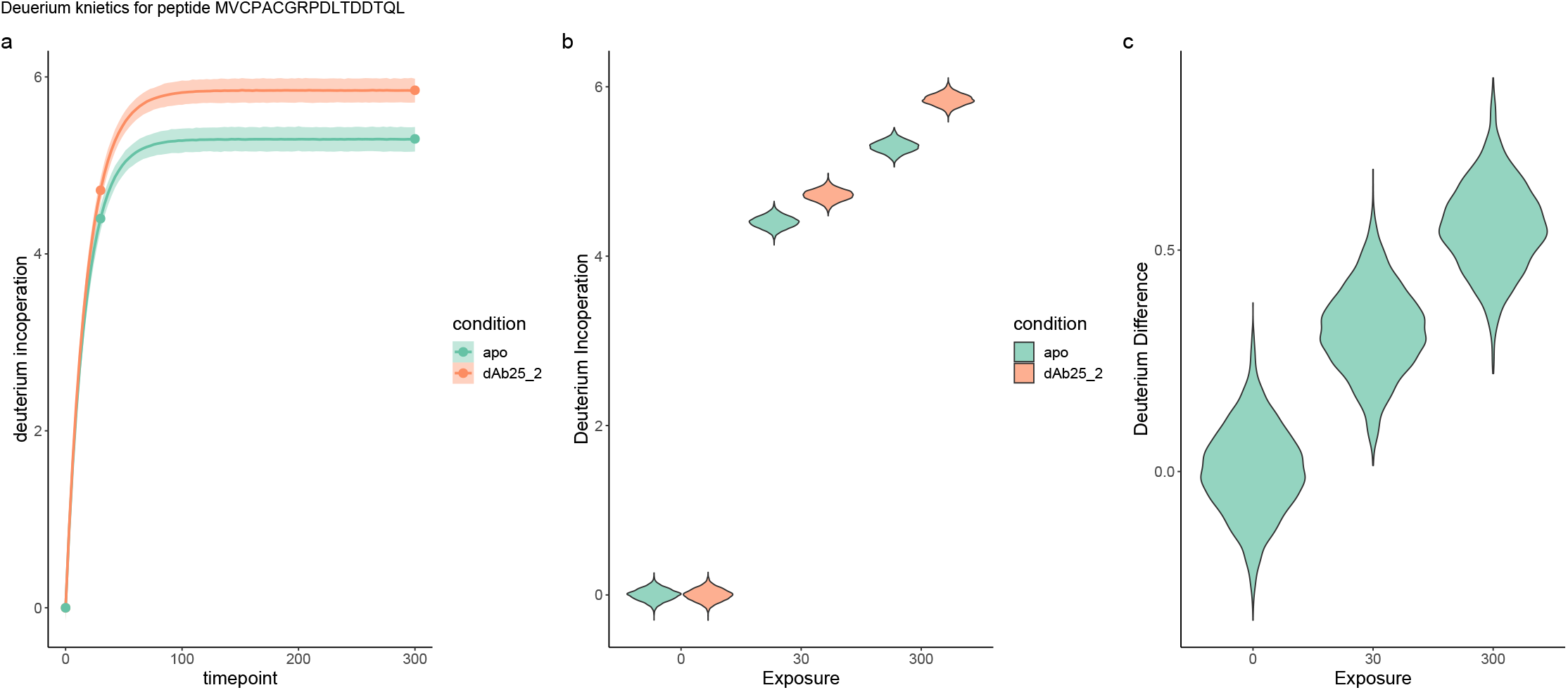
Posterior kinetics for peptide from the Bayesian analysis. (a) The posterior predictive logistic distribution for peptide across the temporal dimension. (b) Posterior predictive distribution of deuterium incorporation for each state. Violin plots represent uncertainty in the deuterium incorporation. (c) Deuterium differences calculated from the posterior predictive distributions. We can clearly see that the probability that the deuterium difference is greater than 0 is close to 1 at 30 and 300 seconds.

#### 2.4.2 HDX-MS analysis of HOIP-RBR-dAb3

We now turn to another epitope mapping task from the HOIP-RBR dataset. Tsai *et al*. (2020) report an X-ray crystal structure of HOIP-RBR-dAB3, where dAb3 forms a dimer (see Figure 6 d). Tsai *et al*. (2020) show that the dAb3 dimer contacts the IBR domain of HOIP from residues on the complementarity determining regions (CDRs). Residues on each monomer form hydrogen bonds with two distinct clusters of residue on HOIP suggesting that the dAb3 dimer has interactions with different sections of the IBR. However, no previous analysis of HDX-MS data on this system could identify any binding epitopes. More specifically, application of the empirical Bayes functional method led to no peptides being identified as having significant changes in deuterium incorporation in this experiment (Crook *et al*., 2022b). Hence, we apply our Bayesian approach to determine whether it could identify these interaction from the HDX-MS data.

**Figure 6:**
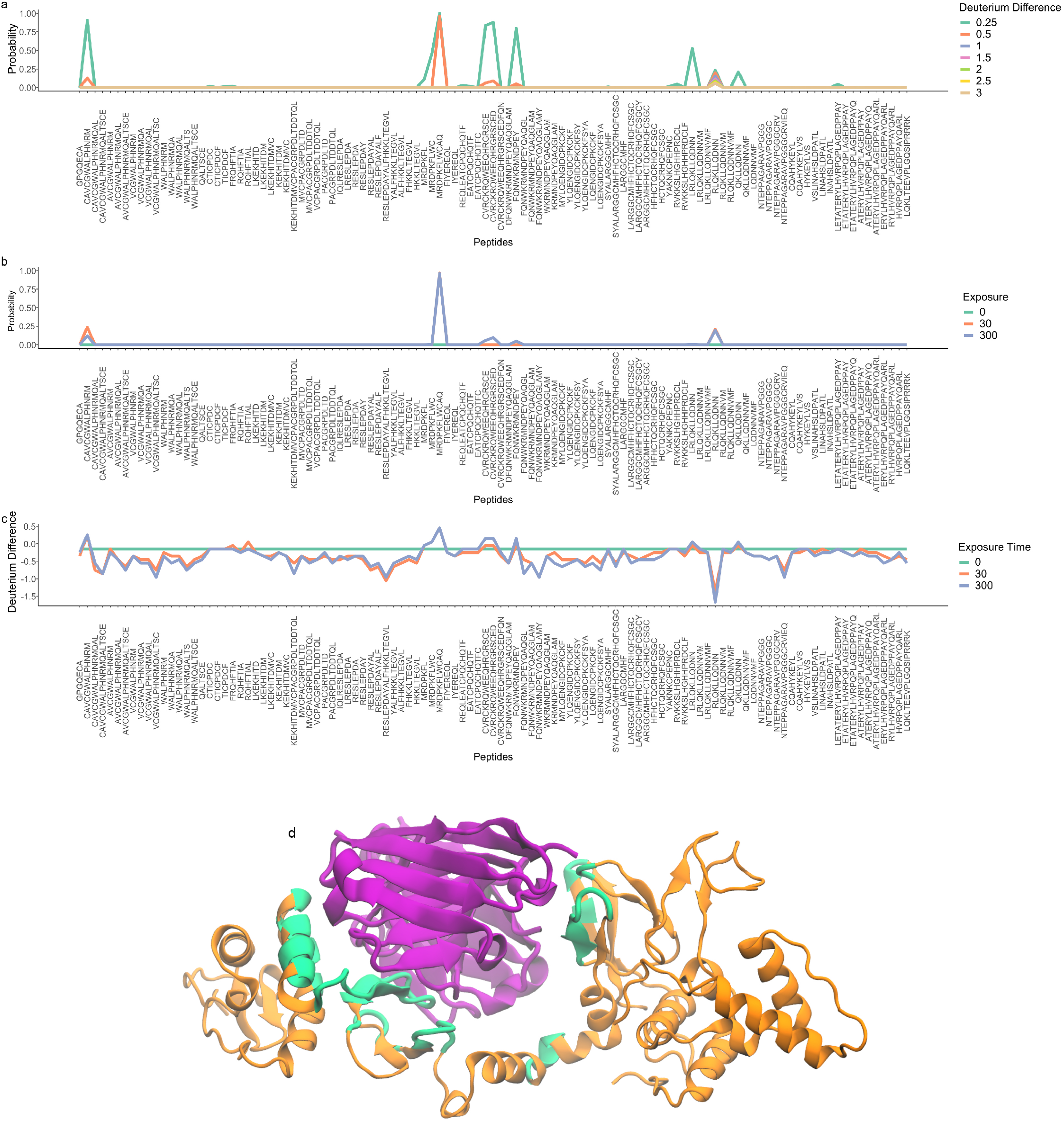
Posterior quantities to identify the epitope in HOIP-RBR-dAb3. (a) The probability that the deuterium incorporation is reduced by a particular effect size in the dAb bound state. (b) The temporal consistency of the computation in (a) with *d* = 0.5. (c) The largest value of *d* such that *P*(Δ*D* > *d*) = 0.95 for different time points plotted as a function of peptide. (d) Cartoon representation of HOIP-RBR-dAB3. Distance analysis of co-complex of HOIP-RBR-dAB3. Residues within 8Å of the dAb3 dimer (violet) are highlighted in green on HOIP-RBR (orange) PDB: 6SC6.

Figure 6 a,b, and c plot the relevant probabilistic calculations for epitope mapping that we previously outlined for HOIP-RBR-dAb25. From visual inspection of these plots, we identify peptides indicative of an epitope. These three peptides include residues that were identified interacting with dAb3 in the crystal structure: Arginine (R792) and Aspartic Acid (D793) (Tsai *et al*., 2020). These residues interact with residues on the CDR2 region of first dAb3 monomer. Confirming that HDX-MS could localise the first binding interface in agreement with the crystal structure. A second small cluster of two peptides shows evidence for small deuterium differences. These two peptides include an Arginine (R827) and Lysine (K829) which were also identified to interact with the CDR3 region of the second dAb3 monomer. Again, demonstrating concordance between our Bayesian analysis of HDX-MS data and the crystal structure. A third potential, but less probable, location for an epitope is in peptide FQNWKRMNDPEY [844 - 855] (see supplementary figure 14 c), though it is not supported by overlapping peptides. Performing distance analysis on the co-crystal structure of HOIP-RBR-dAb3 (PDB: 6SC6), we find 3 interfaces within 8 Å of the dAb3 dimer (Figure 6d). This includes a segment between D766 and E809; another segment between C817 and W832 and finally between R849 and Q858. These three interfaces are concodant with HDX findings and suggest that Bayesian analysis combined with HDX is able to sub-localise interaction sites.

Other peptides which appear to have some changes to their deuterium incorporation are either at low probability or are inconsistent with overlapping peptides, suggesting these are more likely random fluctuations rather than possible binding epitopes. This shows that bespoke statistical approaches for HDX-MS data can provide complementary and consistent evidence to other structural data which was not possible with other statistical methods.

Next, we performed differential solvent accessibility analysis for HOIP-RBR-dAb3. Briefly, we computed the accessible solvent area (ASA) for each residue of the unbound and bound forms of HOIP-RBR (Heinig and Frishman, 2004). We identified two residues with confident differences (see methods and supplementary figure 13). The first is residue D793, which is contained within confident peptides displaying signatures of solvent occlusion. The second is residue R928, which is distal from the hypothesised interaction sites. The relevant peptide from HDX-MS analysis is RVKKSLHGHHPRCVL [917-931] (see supplementary figure 14 d), for which we observed a small but non-zero probability in the perturbation of its HDX kinetics. Examining the crystal structure, we see a rearrangement of the turn on which this residue is localised: *ψ*_[927,928,929]_ = (173, −35, −37) in unbound form and in bound form *ψ*_[927,928,929]_ = (162, −18, 16). The apparent contradiction with HDX-MS analysis, which suggests a small difference, and the large changes observed in the crystal structure arises for two reasons. The first is the signal for HDX-MS analysis is averaged over the residues of the measured peptide. Secondly, the crystal structure is only a single conformation of HOIP-RBR-dAb3, whilst the HDX-MS analysis is again averaged over the ensemble of possible conformations. This highlights the utility of HDX-MS to report on the ensemble of structures and the ability of Bayesian analysis to identify subtle changes. Table 1 summarises the agreement of the co-crystal structure and the HDX-MS data.

**Table 1:**
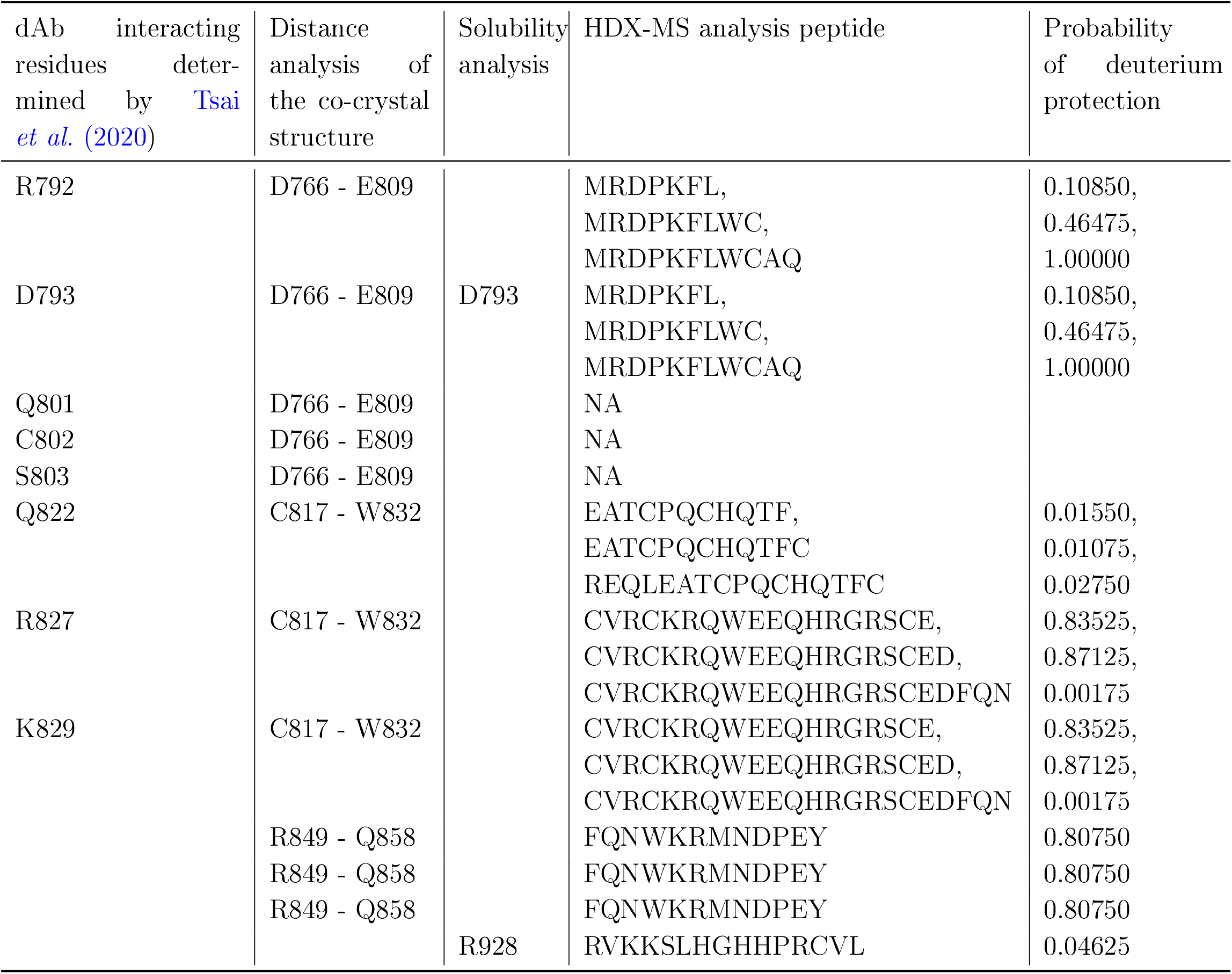
A summary of the correspondence between the Bayesian analysis of HDX-MS data and the co-crystal structure. Distance analysis is performed at eight Å. Solubility analysis is performed as described in the supplement. HDX-MS analysis peptide indicates the measured peptide. NA is used to indicate no peptide with exchangeable residues was measured for that residue.

We further explored the 3 peptides that overlapped with R792 and D793, so that we could carefully examine their kinetics. Time-resolved kinetic plots and violin plots (Figure 7), show that the posterior predictive distributions overlap for peptides MRDPKFL and MRDPKFLWC, however there is a clear suggestion that antibody binding is reducing deuterium incorporation. Furthermore, the posterior predictive distribution for MRDPK-FLWCAQ shows much clearer evidence for a reduction in deuterium incorporation. Due to back-exchange it is unlikely that the HDX-MS measurements are reporting on a known interaction at residue Q801, suggesting that residues C799 or A800 are also important. Examining the co-crystal structure of the dAb3 dimer and HOIP-RBR, we see that the loop containing C799 and A800 are within 8Å of the first dAb3 monomer and so potentially interacting (Figure 6d). We then carefully examined HOIP-RBR-dAb3 (bound HOIP) and HOIP in complex with the ubiquitin-transfer complex (free HOIP), zooming in on residues C799 and A800 (Figure 7j). In bound HOIP, we see that the backbone nitrogen of C799 forms a clear hydrogen bond with the carbonyl from residue F804, and the nearby zinc cation suggests that this loop is structured ((Figure 7j)). Meanwhile, in unbound HOIP, we observe the same residue contacts. Hence, the proximity of the antibody likely stabilises this loop and reduces HDX at C799. In contrast, in both bound and unbound HOIP the amide of A800 is likely bound to water suggesting that it can freely exchange, with no hydrogen bond directly formed with the antibody dimer. This suggests that C799 is a critical residue for this interaction. Hence, the results of our Bayesian analysis on HDX-MS data are consistent with the X-ray crystal structure of HOIP-RBR-dAb3 and complementary information can sub-localise binding epitopes.

**Figure 7:**
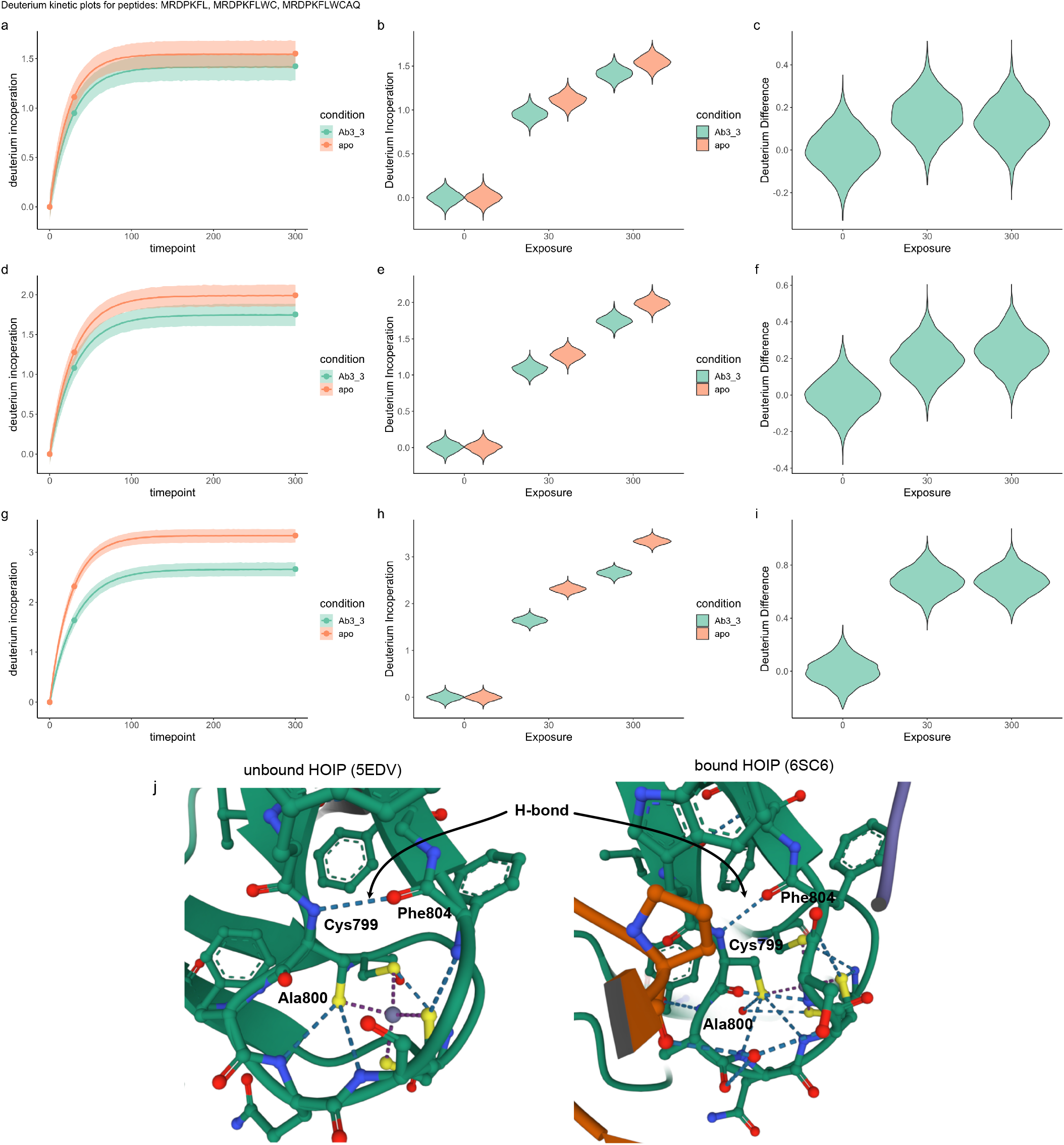
Posterior predictive kinetic plots for three peptides. MRDPKFL is plotted on row 1; MRDPKFLWC on row 2 and MRDPKFLWCAQ on row 3 (a,d,g) The posterior predictive logistic distribution for peptide across the temporal dimension. (b,e,h) Posterior predictive distribution of deuterium incorporation for each state. Violin plots represent uncertainty in the deuterium incorporation. (c,f,i) Deuterium differences calculated from the posterior predictive distributions. Violin plots overlap with zero for exposure times of 30 and 300 seconds. (j) zooms of amides of C799 and A800 on HOIP-RBR in complex with dAb3 dimer (right) and HOIP in complex with the ubiquitin-transfer complex (left). Antigen is coloured in dark green, antibody dimer in brown and purple, elements are coloured using standard colourings. The zinc cation is denoted by a grey sphere and waters are not shown. The corresponding hydrogen bond between C799 and F704 are annotated. Note that A800 does not directly form a hydrogen bond with the dAb3 dimer.

## 3 Discussion

Current statistical approaches applied to or developed for HDX-MS data lack flexibility. For example, t-tests and linear mixed models are only applicable with sufficiently large numbers of replicates. A recently proposed empirical Bayes approach was able to identify difference when the t-test and linear mixed models where not applicable (Crook *et al*., 2022b). However, that approach was still unable to identify difference in some HDX-MS experiments (Crook *et al*., 2022b). To overcome these limitations, we developed a Bayesian approach to modelling HDX-MS data. Our approach combines a likelihood model that captures the time-dependent kinetics of HDX data, either as a logistic or Weibull model. We place priors on the parameters of these likelihood models, which rules-out unrealistic parameter values and allows us to shrink residuals towards zero. We sample from the posterior distribution of this model using a variant of Hamiltonian Monte Carlo (HMC) (Hoffman and Gelman, 2014; Betancourt, 2017). This posterior distribution allows us to quantify uncertainty in the parameters of our model or, more generally, any posterior quantity that we are interested in (Gelman *et al*., 2013).

Through careful simulations, we show that our Bayesian model correctly identifies true perturbations and has similarly high performance when compared to the previous empirical Bayes approach. We also demonstrate that our model is correctly calibrated by examining the Brier score (Gneiting and Raftery, 2007). We then use simulations and a structural spike-in experiment to demonstrate the power of Bayesian model selection (Gelfand and Dey, 1994). Here, we use the Bayesian tool-kit to demonstrate that EX1 kinetics are modelled better by Weibull model compared with a logistic model. Furthermore, we rule out a more complex random plateau model. Then, we showed that our Bayesian approach does not unnecessarily inflate false positives.

We then turn to an epitope mapping experiment for HOIP-RBR, an E3 ubiquitin ligase, involved in immune signalling (Kirisako *et al*., 2006). We first consider HOIP-RBR-dAb25, to demonstrate a number of new computations and visualisation. Our analysis suggests that a region of the IBR contains the binding epitope, by identifying 7 key peptides in this region with reduced HDX. This more than triples the evidence for an epitope in this region compared with previous analysis (Tsai *et al*., 2020; Crook *et al*., 2022b). Our analysis also reaffirms the idea that this antibody holds HOIP-RBR in a more open conformation, as most peptides have a tendency to increase their deuterium incorporation in the bound state. We proceed to consider HOIP-RBR-dAb3 because previously it has not been possible to identify an epitope from this HDX data (Tsai *et al*., 2020; Crook *et al*., 2022b). However, our Bayesian analysis suggests three possible interfaces on the IBR that could be the binding epitope. All these regions are consistent with distance analysis on the co-crystal structure of HOIP-RBR-dAb3. Using this crystal structure, we sought to further interrogate the HDX kinetics. We suggest that the dAb3 dimer likely stabilises the interaction network of residue C799. Our findings demonstrate a Bayesian analysis can identify important binding epitopes from HDX-MS data that could not be found with other approaches. This methodology presents a powerful alternative approach to the statistical analysis of HDX-MS, allowing us to quantify the uncertainties in our analysis.

## 4 Methods

### 4.1 Preliminaries

In hydrogen deuterium exchange mass spectrometry, we observe isotopic distributions for *i* = 1,…, *n* peptides at different exposure times *t*_1_, ‥, *t*_*m*_ to heavy water (D_2_O). The isotope distributions are a set of 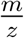-Intensity pairs reporting the relative intensities of each peptide isotope. These isotope distributions are frequently summarised by the intensity weighted mean of the 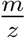, which we write as 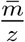 and is referred to as the centroid. Since deuterium is heavier than hydrogen, deuterium interoperation leads to positive shifts in 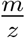 and we monitor this change over time and with respect to the state. In most scenarios data are replicated, so we observe replicates *r* = 1,…, *R* and, potentially, a number of conditions/covariates denoted *c =* 1,‥, *C*. The observations are then

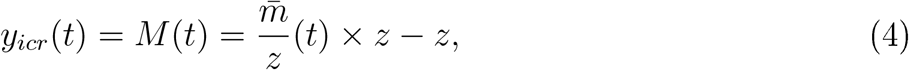

where *z* denotes the charge of the precursor ion. It is typical to normalise such that mass *M*(0) = 0.

### 4.2 Bayes’ theorem and hypothesis testing

Here, we summarise Bayesian inference and hypothesis testing (Gelfand and Dey, 1994; Gelman *et al*., 2013). We first note that multiplicity is implicitly controlled via prior distributions on the parameters, as well as explicitly via the prior model probabilities. Prior information on the parameters imparts a number of advantages, including shrinkage of residuals towards 0, regularisation of parameters and more stable algorithmic inference. Furthermore, a Bayesian analysis allows us to sample from the posterior distribution of quantities of interest and using this probability distribution is central to quantifying uncertainty.

Bayesian inference requires several quantities to be specified. The first is a statistical model ℳ, with parameters *θ*, of the observed data *y*. After specifying a prior distribution for the parameters, denoted *p*(*θ*|ℳ), and given observed data *y*, Bayes’ theorem tells us we can update the prior distribution to obtain the posterior distribution using the following formula:

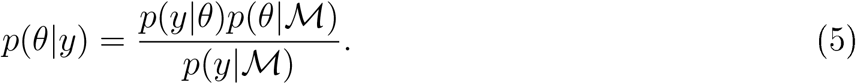

The quantity *p*(*y*|ℳ) is referred to as the marginal likelihood, and is obtained by marginalising *θ*:

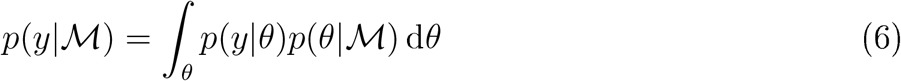

The task of hypothesis testing can be reformulated as a model selection problem. We write ℳ_0_ to denote the model associated with the null hypothesis; whilst the alternative hypothesis is associated to model ℳ_1_. Thus, hypothesis testing can be phrased as selecting between two competing models.

To perform model selection, we are interested in the posterior model probability, given the data (Berger and Molina, 2005):

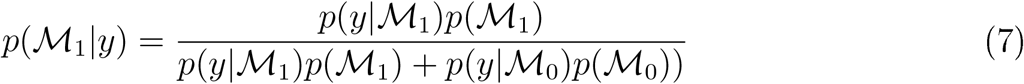

The relative plausibility of two model is quantified through the posterior odds, which is the prior odds multiplied by the Bayes factor (Kass and Raftery, 1995):

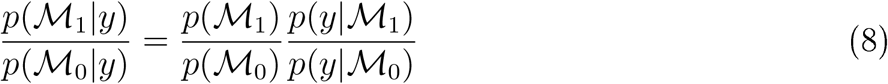

These quantities are challenging to compute because of the marginal likelihoods required. Note that the prior on the parameters penalises additional model complexity. Given that the marginal likelihood is only analytically available for relatively simple models, we approximate it using Bridge sampling (see the following section) (Meng and Wong, 1996; Meng and Schilling, 2002). Though a number of other methods are available such as path sampling (Gelman and Meng, 1998), importance sampling (Robert and Wraith, 2009), harmonic mean sampling (Gelman and Meng, 1998), nested sampling (Skilling *et al*., 2006; Chopin and Robert, 2010) and Laplace’s approximation (Lewis and Raftery, 1997).

Finally, we assume that the null model is more probable than the alternative, controlling multiplicity (Scott and Berger, 2006). Thus, we set *p*(ℳ_0_) = 0.95 and *p*(ℳ_1_) = 0.05.

### 4.3 Bridge sampling for marginal likelihoods

We provide brief details on bridge sampling to compute the marginal likelihood (Meng and Wong, 1996; Meng and Schilling, 2002). We first observe the following identity:

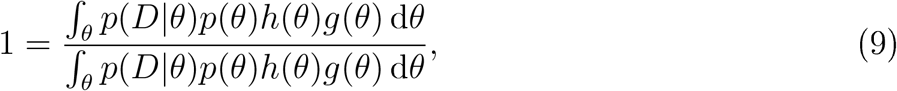

where *g*(*θ*) is the proposal distribution and the bridge function is denoted by *h*(*θ*). Then multiplication of both side by *p*(*D*), we obtain

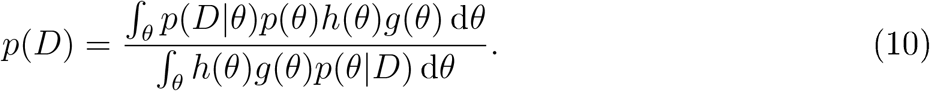

The numerator can be written as an expectation with respect to *g*(*θ*), whilst the denominator is an expectation with respect to the posterior *p*(*θ*|*D*). Hence, we observe that

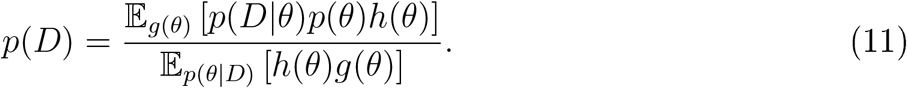

This allows to approximate the marginal likelihood using the following estimator

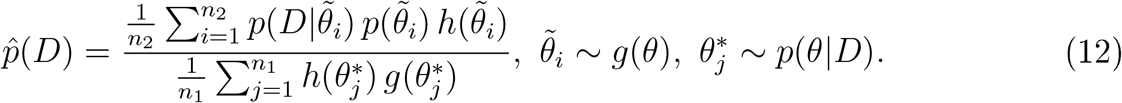

The important observations are that we need samples from the proposal distribution *g*(*θ*) and the posterior *p*(*D*|*θ*). The samples from the posterior are simply obtained from the MCMC algorithm used in model inference. For the proposal distribution to have good empirical performance it should have the same support as the posterior and so a normal proposal with moments matched to the posterior distribution is suitable. We have yet to mention how to decide on the bridge function *h*(*θ*). The so-called optimal bridge function takes the following form (Meng and Schilling, 2002):

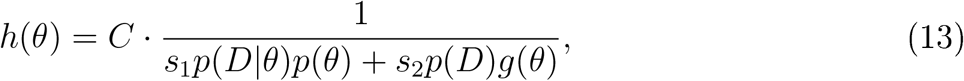

where *s*_1_ = *n*_1_/(*n*_2_ + *n*_1_) and *s*_2_ = *n*_2_/(*n*_1_ + *n*_2_). The constant *C* is irrelevant because in cancels in the estimator ratio. The bridge function is optimal in the sense that in minimises relative mean-squared error. However, a clear issue with this choice of bridge function is that it depends on the quantity we are trying to estimate *p*(*D*). The resolution is to make an initial guess for the marginal likelihood *p*^(0)^(*D*) and iteratively update accordingly. The update equation is given by

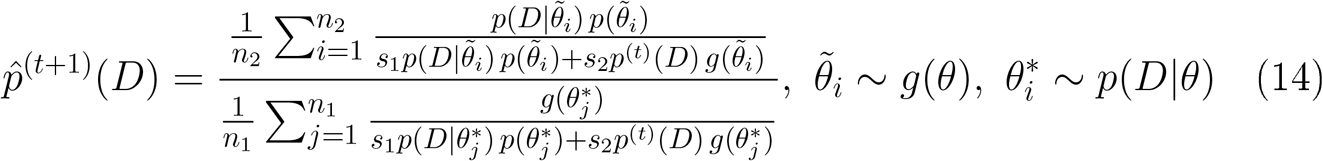

The above iterative estimator demonstrates why bridge sampling is robust to the tail behaviour of proposal distribution relative to the posterior distribution - a property not held by other methods such as importance sampling.

### 4.4 Prior and posterior predictive checks

In this section, we summarise prior and posterior predictive checks which allow us to build high quality generative and predictive models (Box, 1980; Gabry *et al*., 2019; Betancourt, 2021). Given a likelihood and prior, we can simulate data 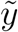 (Gelman *et al*., 2017). First, sample the parameters of the likelihood from the prior and then given these parameters sample data from the model:

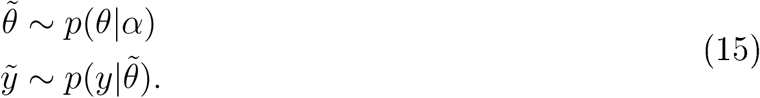

This leads us to define the *prior predictive distribution*:

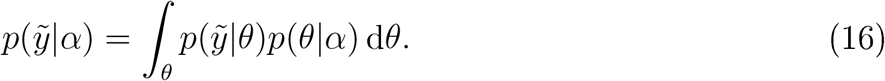

See Gelman *et al*. (2017) for advantages of working with the prior predictive distributions. To define the posterior predictive distribution, we can then sample new data by first sampling parameters from the posterior distribution and then again sampling from the likelihood:

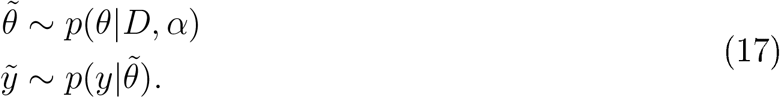

This leads to the posterior predictive distribution:

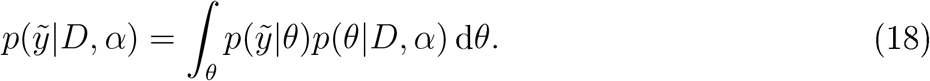

Once we have data 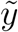 generated from a predictive distribution we can generated a summary of this data 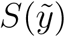. We can then visually compare the summary of an ensemble of simulated datasets 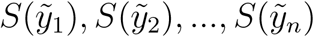, with the summary of the observed data *S*(*y*) to identify model deficiencies.

### 4.5 Out-of-sample predictive performance

Another approach to evaluate the quality of a Bayesian model is to examine the out-of-sample predictive accuracy from the fitted model. We can employ (approximate) leave-one-out cross validation (LOO-CV) with log predictive density as the utility function (Vehtari *et al*., 2017):

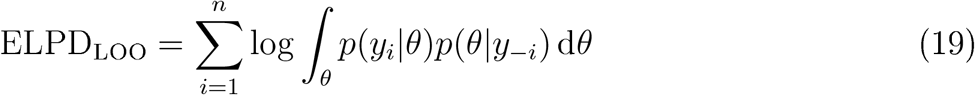

The above equation is the leave-one-out predictive density given the observed data with the *i*^*th*^ observation removed, summed over the observations. This quantity is estimated using Pareto smoothed importance sampling (PSIS) (Vehtari *et al*., 2015).

### 4.6 Modelling

Here, we state the proposed models considered in this manuscript. This includes a logistic model, a Weibull model and a random-effects model. In each of these cases, we consider a model ℳ_0_ in which a single function is posited and blinded to any contexts, covariates or treatment effects. Meanwhile, we also consider allowing the parameters of the model, ℳ_1_, to be context-specific.

For the *logistic* model, we assume

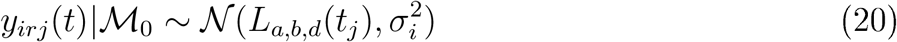

and

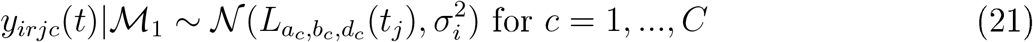

where *c* denotes the context. The mean logistic model is given by:

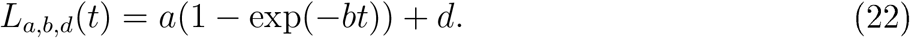

To complete the specification of the model, we need to specify priors. The first parameter is *d* which can be interpreted as an intercept, which should be symmetric and concentrate around 0. The parameter *a* represents the plateau of the model (dominates for large values of *t*). This parameter is also required to be positive. The parameter *b* is a shape parameter and also is required to be positive. The standard deviation *σ* also must be positive, furthermore we wish to penalise this parameter to avoid concentrating on models which are completely explained by noise. These considerations lead to the following hierarchical structure:

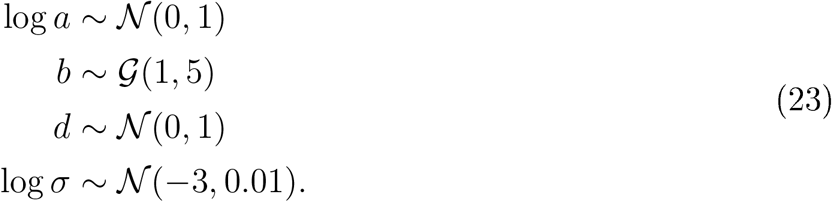

These priors were chosen using iterative prior predictive checks and the quality of posterior inference was examined using posterior predictive checks (see supplement).

We also consider a *Weibull* model for HDX-MS data, which allows more flexibility in the temporal kinetics. As for the logistic model, we posit

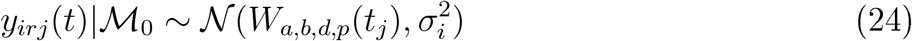

and

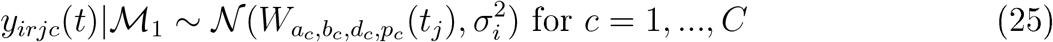

where *c* denotes the context. The mean logistic model is given by:

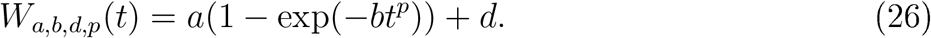

The prior construction is the same for the logistic model, but in addition we propose:

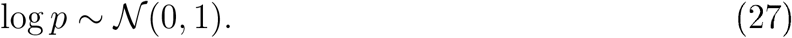

Finally, we consider a functional random-effects Weibull model by allowing random plateaus using a replicate level grouping nested within condition. That is, the parameter *a* is modelled as follows, denoting *r* for replicate

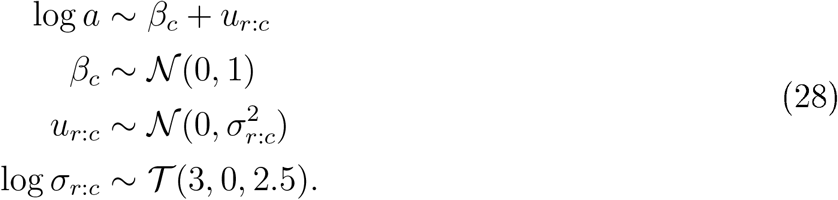

### 4.7 Inference and implementation

Model and parameter inference is performed using Markov-chain Monte-Carlo (MCMC); in particular, we use the No U-Turn Sampler (NUTS) a variant of Hamiltonian Monte-Carlo, as implement in the probabilistic programming language stan (Betancourt, 2017; Carpenter *et al*., 2017).

### 4.8 Previous approaches

We compare to an empirical Bayes functional modelling approach (Crook *et al*., 2022b). Briefly, Weibull or logistic functional models are fitted to the time-dependent HDX kinetics. By fitting a null model that is blinded to the covariates and an alternative hypothesis that uses independent models for each context, we are able to compute the residual sum of squares for each model. From this, we can compute an F-statistic, with the appropriate degrees of freedom. To borrow power, an empirical Bayes method is applied allowing the variances to be shrunk toward a pooled variance. This stabilises variance estimates and improves power by computing a *moderated* F-statistic. *p*-values are then computed from the appropriate F-distribution and then corrected for multiple testing using the Benjamini-Hochberg procedure.

### 4.9 Performance metrics

To compare the Bayesian approach with the empirical Bayes functional method, we use the area-under-curve of receiver operating characteristics (AUCROC). That is the area under a curve by plotting the true positive rate (TPR) against the false positive rate (FPR). AUCROC of 1 indicates perfect performance, whilst 0.5 denotes random performance. To assess the calibration of probabilities from the Bayesian approach, we compute the Brier score. A Brier score of 0 indicates probabilities that are perfectly calibrated and a Brier score of 1 indicates probabilities that display no calibration (Gneiting and Raftery, 2007).

### 4.10 Simulation study

This section describes our proposed simulation study, as in (Crook *et al*., 2022b). We begin by sampling, uniformly at random, the length of the peptide between 5 and 25. The sampled number is the number of amino acids in the peptide and we sample that number of amino acids from the 20 canonical amino acids with replacement. We then define time points at which to obtain data: *T =* {*t*_1_, …, *t*_*m*_}, with *t*_1_ = 0 and *t*_*i*_ < *t*_*j*_ for *i* < *j*. For time *t*_1_, we simulate the undeuterated isotope distribution using a binomial modal. For a subsequent time point *t*_*i*_ we sample the percentage incorporation by first sampling from a *m* − 1-variate Dirichlet distribution with concentration parameter *α*, where *α*_*i*_ = 20/(*i* − 1). From this we obtain a vector *π* which sums to 1. We use the cumulative distribution of *π* as the schedule of incorporations. That is the incorporation at 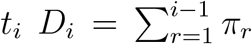 for *i* > 1. This ensure that incorporation is non-decreasing in time. To simulate the effect of a condition, for each time point, we sample an indicator 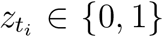 such that the 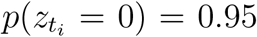. If 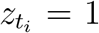, then we re-sample the incorporation amount and continue on the simulation process. This ensures that roughly 95% of the scenarios have no effect with respect to the condition. A binomial model is used to generate deuterated spectra, where the exchangeable hydrogens are randomly replaced with deuterium according to the incorporation percentage. The isotope distribution simulations are repeated *R* times to allow for replicates. Centroids summarising the average peptide mass are then computed from the isotope distribution. In all cases we simulate 100 measured peptides. We perform simulation scenarios as follows:

- (Scenario A) 4 time points, 3 replicates and 2 conditions
- (Scenario B) 4 time points, 2 replicates and 2 conditions
- (Scenario C) 5 time points, 2 replicates and 2 conditions

### 4.11 Differential Solubility Analysis

To perform differential solvent accessibility analysis, we computed the accessible solvent area (ASA) for each residue of the unbound and bound forms of HOIP-RBR. We then took the square-root of these values so that the results were on the linear scale and computed the difference. Finally, results were converted to *z*-scores. Significant differences in ASA were identified by computing the local *false discovery rate* (fdr) (Efron, 2004). We recall that the fdr is the probability that there is no change in the observed differences, for a given test statistic; that is, fdr(*z*_*i*_) = *P*(*z*_*i*_ = 0|*Z* ≤ *z*_*i*_) A threshold of 0.01 was set to declare a difference.

## 5 Acknowledgements

OMC acknowledges funding from GSK, a Todd-Bird Junior Research Fellowship from New College Oxford and the EPSRC (EP/R511742/1). CWC is an employee of GSK.

## 6 Data availability

Data to reproduce the figures is provided in the supplementary material. Experimental data are available from the original manuscripts. Monte-Carlo data is available from (10.5281/zcn-odo.6855078)

## 7 Code availability

Stan files implementing the methods are provided as part of the supplementary material.

## 8 Supplementary material

### 8.1 Prior and posterior predictive checks

**Figure 8:**
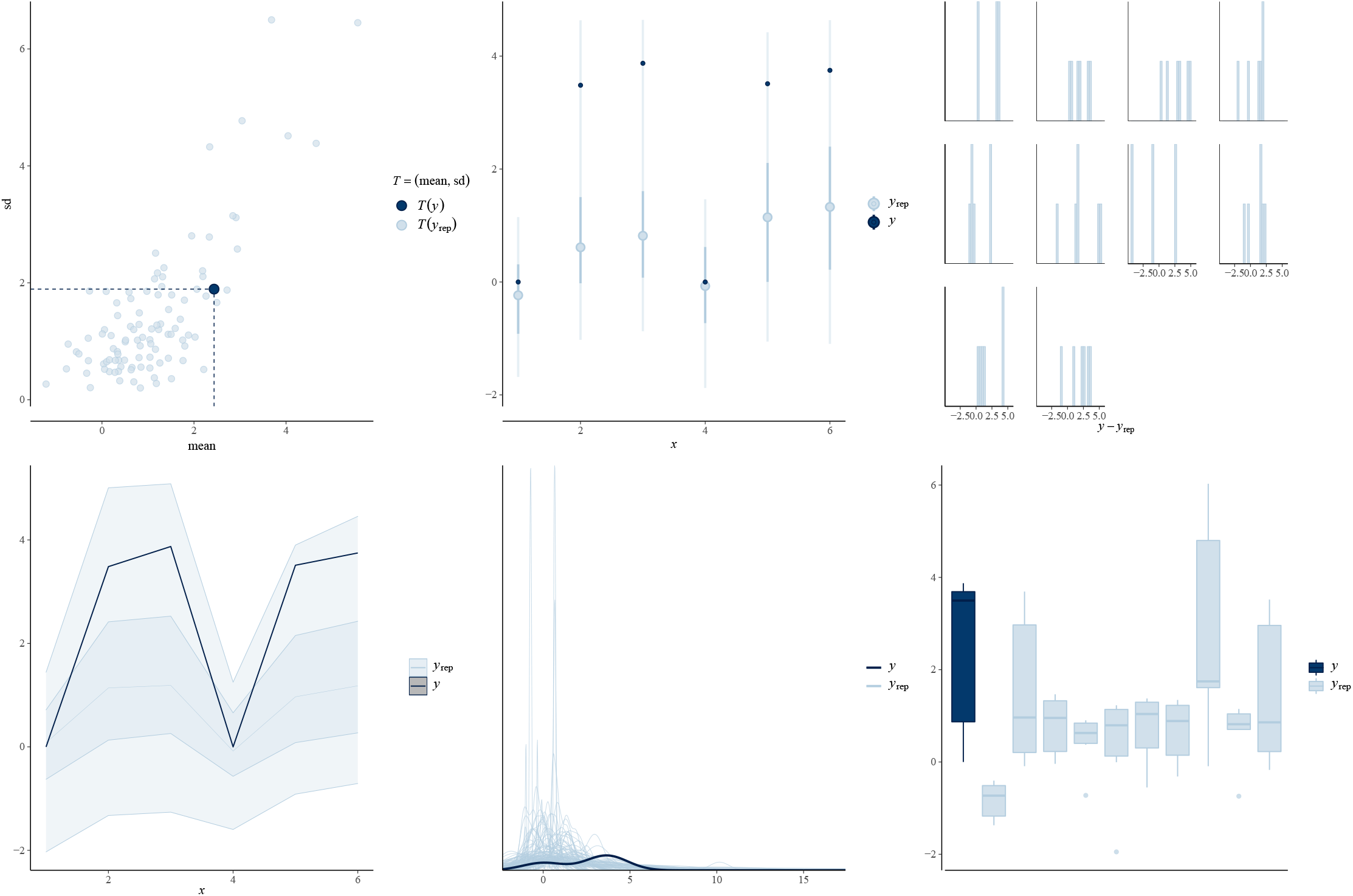
Example prior predictive checks. Each panel shows a prior predictive check for a different summary statistics. We simulate from the prior predictive distribution denoted as *y*_*rep*_ and compare to the observed data *y*. We see that the prior predictive is diffuse but correctly located

**Figure 9:**
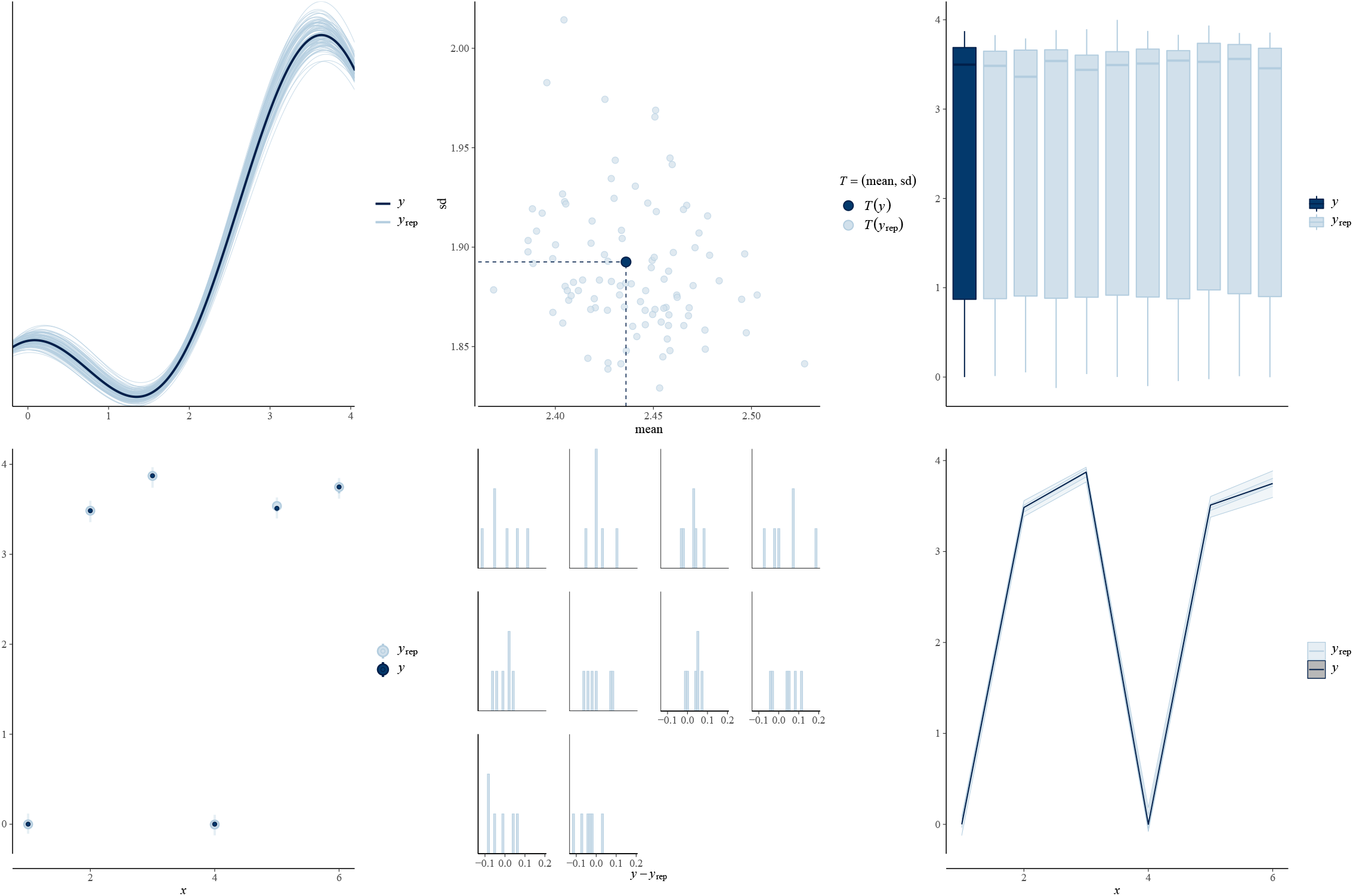
Example posterior predictive checks. Each panel shows a posterior predictive check for a different summary statistics. We simulate from the posterior predictive distribution denoted as *y*_*rep*_ and compare to the observed data *y*. We see that the posterior predictive is concentrated and correctly located

### 8.2 Convergence analysis

**Figure 10:**
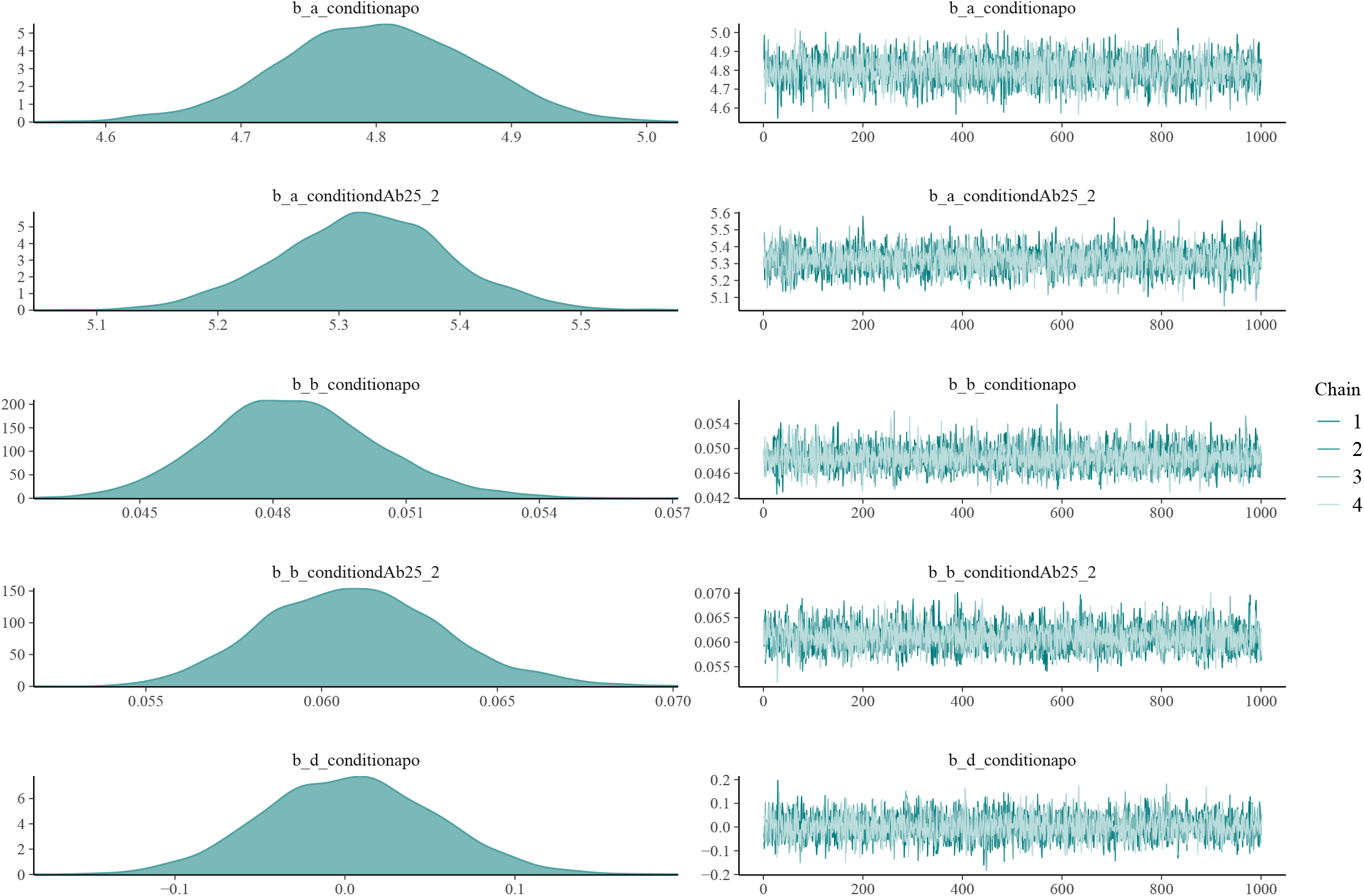
Example trace plots. Trace plots for example parameters from MCMC runs, convergence is clearly achieved

**Figure 11:**
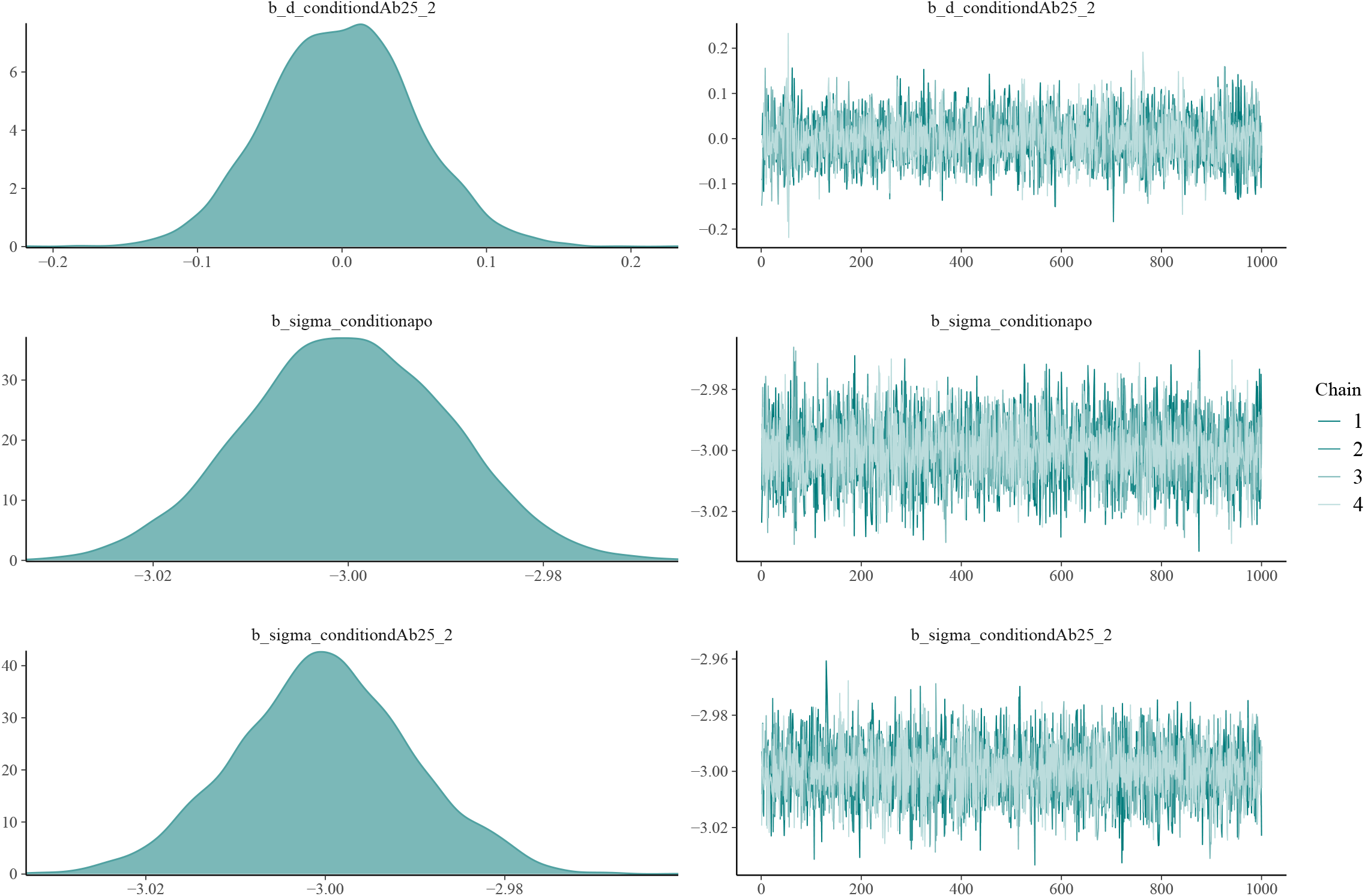
Example trace plots. Trace plots for example parameters from MCMC runs, convergence is clearly achieved

### 8.3 Control of error rates in null experiments

To demonstrate that our Bayesian analysis controls false discoveries, we perform a permutation experiment, using the experiment on MBP generated in seven replicates introduced in the main text. The seven MBP samples without any structural variant can be used as a null experiment by partitioning the replicates *falsely* into two conditions. That is three of the samples are labelled condition *A* and four samples are labelled condition *B*, arbitrarily. We randomly permute the samples labelled *A* and *B*, five times. We then computed the posterior probability that each peptide is perturbed (alternative model). For each permutation this is visualised in a histogram of all the probabilities. We see that the posterior probability is never above 0.05 suggesting excellent control of error rates; that is, we never give confident support to the wrong model. We also check that the probabilities are calibrated by computing the Brier score for each permutation experiment. We plot the Brier scores as a boxplot and see that they are essentially 0, indicating good calibration.

**Figure 12:**
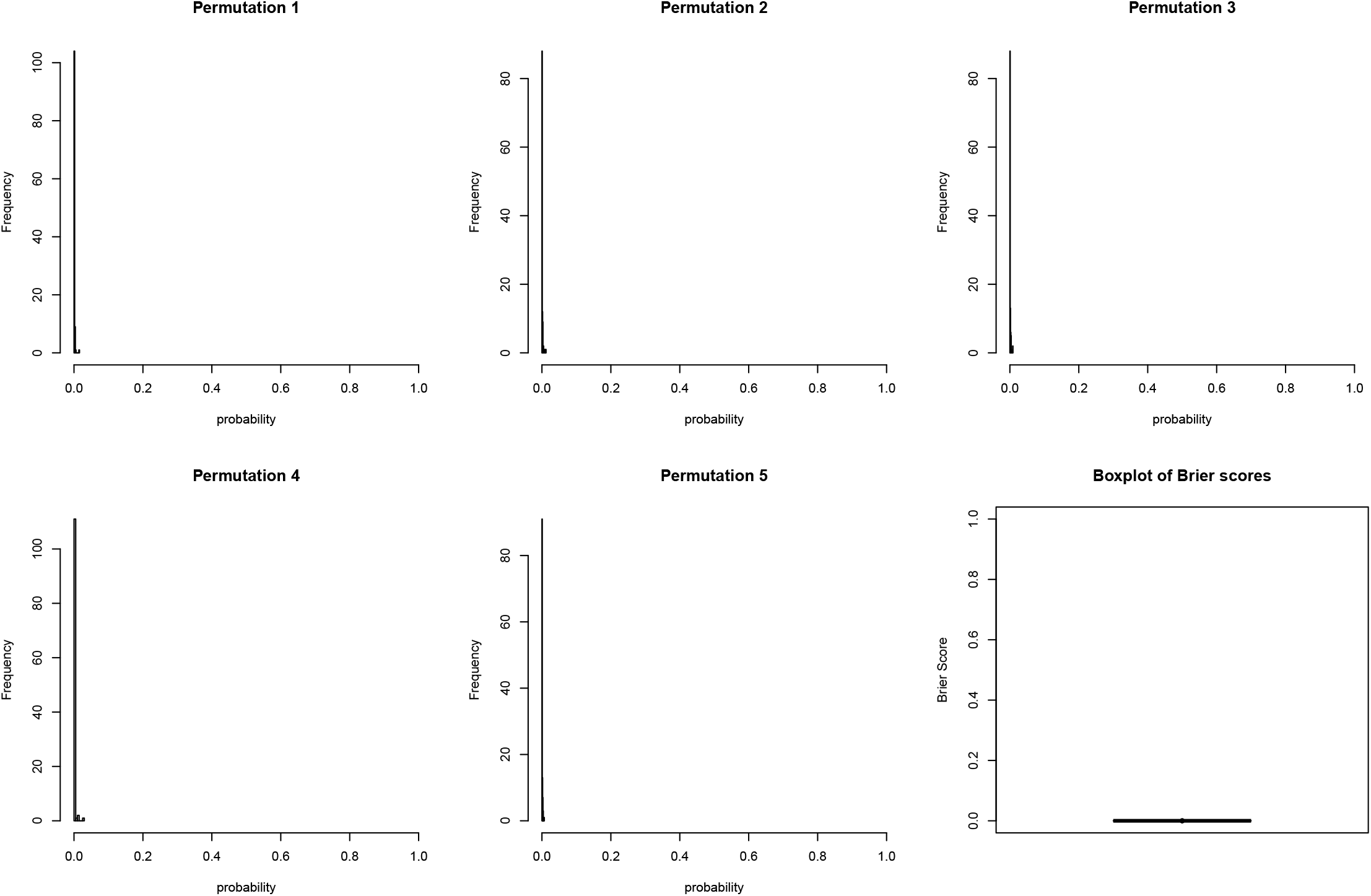
Bayesian analysis controls error rates in null experiments. (1-5) Histograms for the computed posterior probability of the alternative model. The alternative model is clearly never strongly supported (6) The histogram of Brier scores for each null permutation experiments. The values are close to 0

### 8.4 Differential solvent accessibility analysis

**Figure 13:**
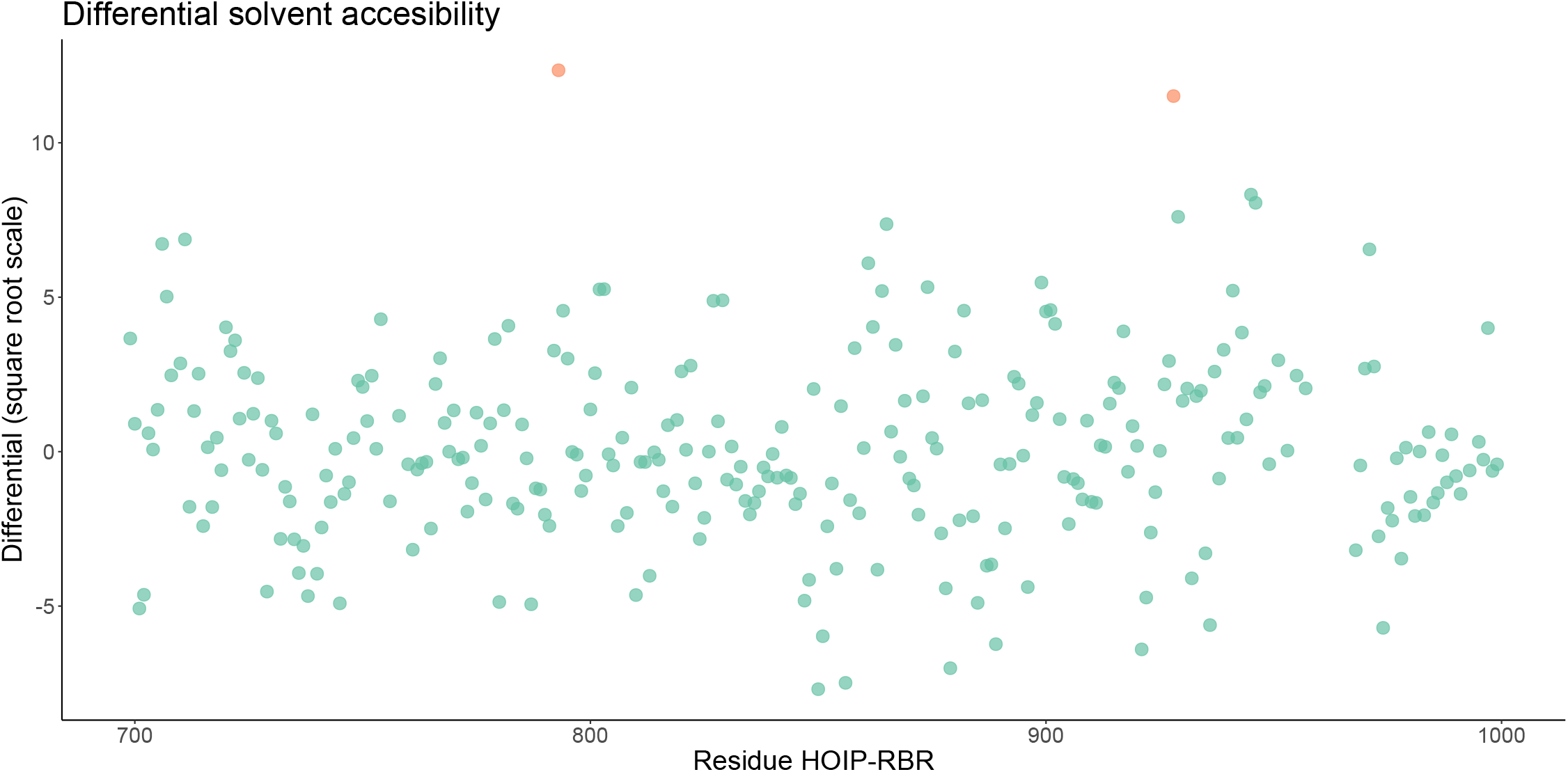
Differential solvent accessibility analysis. Differential solvent accessibility analysis of HOIP-RBR in bound (6SC6) and unbound (5EDV) forms. The difference in solvent accessibility is plotted on the squart-root scale against the HOIP-RBR residue number. Significantly difference are identified by computing the local *false discovery rate* (FDR) and highlighted in orange.

### 8.5 HOIP-RBR structure and peptides

Here, we plot the structure of HOIP-RBR and highlight peptides referenced in the main manuscript

**Figure 14:**
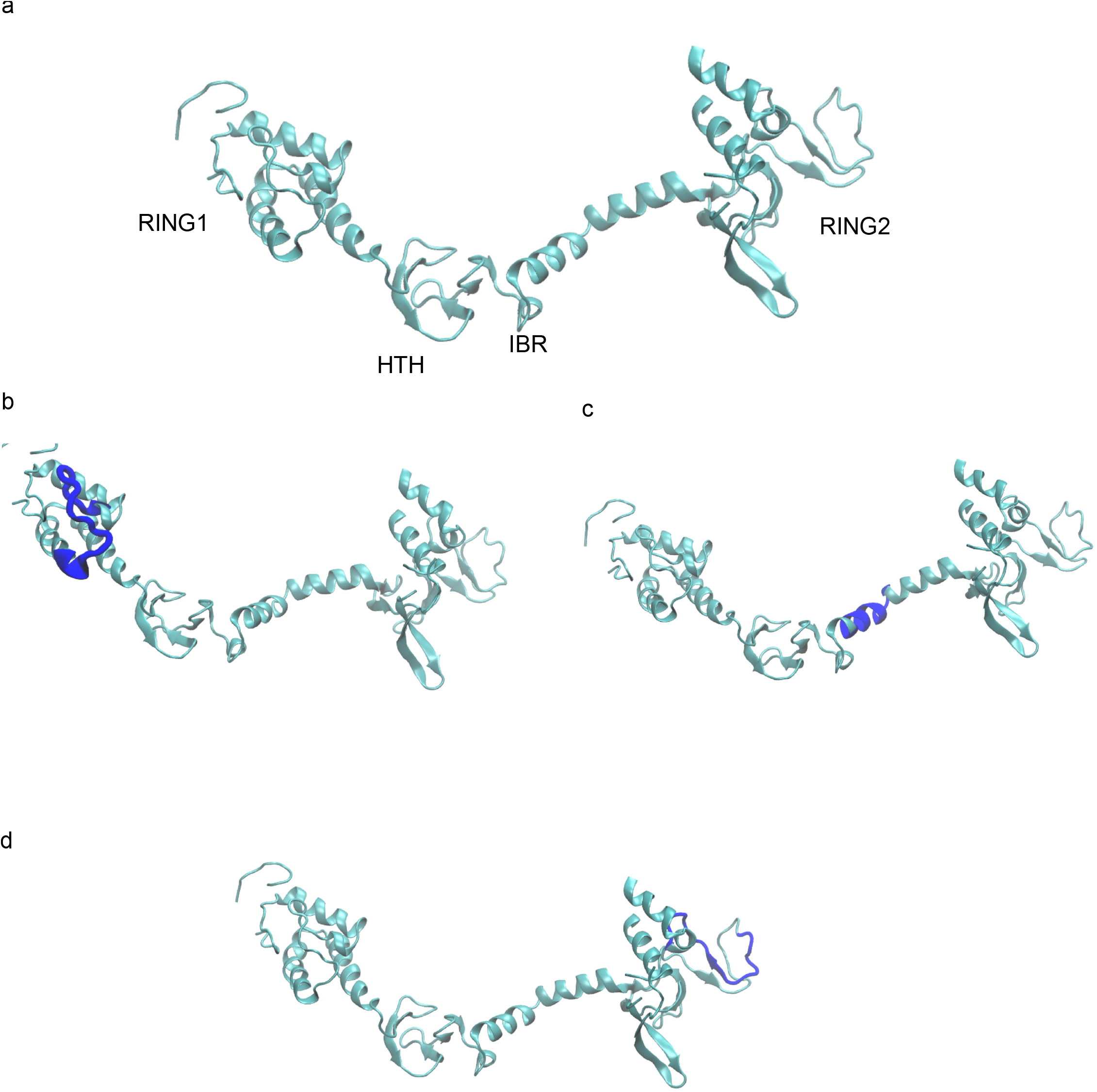
HOIP-RBR structures. (a) HOIP-RBR structure (pdb: 5EDV) (b) peptide 742-758 (c) peptide 844-855 (d) peptide 917-931

